# *Foxc1* and *Foxc2* function in osteochondral progenitors for the progression through chondrocyte hypertrophy and mineralization of the primary ossification center

**DOI:** 10.1101/2023.04.26.538325

**Authors:** Asra Almubarak, Qiuwan Zhang, Cheng-Hai Zhang, Andrew B. Lassar, Tsutomu Kume, Fred B Berry

## Abstract

The forkhead box transcription factor genes *Foxc1* and *Foxc2* are expressed in the condensing mesenchyme of the developing skeleton prior to the onset of chondrocyte differentiation. To determine the roles of these transcription factors in limb development we deleted both *Foxc1* and *Foxc2* in lateral plate mesoderm using the Prx1-cre mouse line. Resulting compound homozygous mice died shortly after birth with exencephaly, and malformations to this sternum and limb skeleton. Notably distal limb structures were preferentially affected, with the autopods displaying reduced or absent mineralization. The radius and tibia bowed and the ulna and fibula were reduced to an unmineralized rudimentary structure. Molecular analysis revealed reduced expression of Ihh leading to reduced proliferation and delayed chondrocyte hypertrophy at E14.5. At later ages, Prx1-cre;Foxc1^Δ/Δ^;Foxc2 ^Δ/Δ^ embryos exhibited restored Ihh expression and an expanded COLX-positive hypertrophic chondrocyte region, indicating a delayed exit and impaired remodeling of the hypertrophic chondrocytes. Osteoblast differentiation and mineralization were disrupted at the osteochondral junction and in the primary ossification center (POC). Levels of OSTEOPONTIN were elevated in the POC of compound homozygous mutants, while expression of Phex was reduced, indicating that impaired OPN processing by PHEX may underlie the mineralization defect we observe. Together our findings suggest that Foxc1 and Foxc2 act at different stages of endochondral ossification. Initially these genes act during the onset of chondrogenesis leading to the formation of hypertrophic chondrocytes. At later stages Foxc1 and Foxc2 are required for remodeling of HC and for Phex expression required for mineralization of the POC.

## Introduction

In mammals, most bones such as the limb skeleton, the vertebrae, ribs and parts of the skull are formed through endochondral ossification (Kozhemyakina et al., 2015). In this process, a cartilaginous template is first formed by chondrocytes to grow and shape the bone before being replaced by bone-forming osteoblasts. In the limb, this event initiates as mesenchymal progenitors from the lateral plate mesoderm condenses and these cells differentiate into chondrocytes (Tickle and Towers, 2020). Once formed, chondrocytes will progress through a series of stepwise differentiation events to form a layered, organized structure known as the growth plate (Yeung Tsang et al., 2014). At the distal end of the growth plate, known as the resting zone, immature chondrocytes transition to a highly proliferative state, and stack together to form the columnar zone of the growth plate. These columnar chondrocytes will then exit from the cell cycle to become pre-hypertrophic chondrocytes which express *Indian Hedgehog (Ihh)* that regulates proliferation and differentiation of chondrocytes and osteoblast formation (Karp et al., 2000; Vortkamp et al., 1996). Prehypertrophic chondrocytes enlarge in volume to become hypertrophic chondrocytes (HC) that form the interface between bone and cartilage and regulate much of the endochondral ossification process. HC express Matrix Metalloprotein (MMP) 9 and 13 which degrades the chondrocyte extracellular matrix (ECM) and along with chondroclasts and osteoclasts, help in the formation of the primary ossification center (POC). HCs produce vascular endothelial growth factor (VEGF) that promotes the invasion of blood vessels that invade the POC (Gerber et al., 1999; Zelzer et al., 2004). Bone-forming osteoblasts from the surrounding perichondrium/periosteum migrate along with blood vessels into POC. In addition, a subset of the osteoblasts that populate the POC are also derived from HC (Yang et al., 2014).

The SRY-Box Transcription Factor 9 *(Sox9)* is an essential regulator of chondrogenesis and endochondral ossification. Mutations to SOX9 causes Campomelic Dysplasia (OMIM#114290) an autosomal dominant disorder that results in shortened, bowed limbs and diminished rib cage formation often leading to perinatal death (Foster et al., 1994). Analysis of SOX9 function has led to important understanding of chondrogenesis and endochondral ossification (Liu et al., 2017). Sox9 is expressed in condensing mesenchymal cells and initiates chondrogenesis in the developing bone anlage (Bi et al., 1999). SOX9 regulates expression of Collagen 2 alpha 1, an abundant extracellular matrix protein of chondrocytes (Bell et al., 1997), as well as other factors that control progression through chondrogenesis (Liu et al., 2017). Although Sox9 function is important for chondrogenesis to occur, it is not essential as rudimentary chondrocytes can form in its absence, and thus, other factors must regulate this process, either working with SOX9 or independently of it (Liu et al., 2018).

The transcription factors Forkhead Box C1 and C2 (FOXC1 and FOXC2) are important regulators of skeletal development (Almubarak et al., 2021; Kume et al., 1998; Winnier et al., 1997; Yoshida et al., 2015). These factors display overlapping expression in skeletal progenitors of the limb and the vertebrae (Almubarak et al., 2021; Hiemisch et al., 1998). Mice with homozygous null mutations in either *Foxc1* and *Foxc2* display malformations in the axial skeleton (the skull, vertebral column, and rib cage). In contrast, the bones in the limbs or appendicular skeleton are less affected (Hong et al., 1999; Kume et al., 1998; Winnier et al., 1997). Compound *Foxc1^-/-^; Foxc2^-/-^* mice die around Embryonic (E) day 9 due to failure in cardiovascular development, and before any skeletal structures are formed, preventing any analysis of the possible association between the two transcription factors in endochondral ossification to be studied (Kume et al., 2001). To address potential compensation between Foxc1 and Foxc2, we created a conditional mouse model that deleted both genes in the chondrocyte lineage. These Col2-cre;*Foxc1^Δ/Δ^;Foxc2^Δ/Δ^*mutants displayed impaired chondrocyte differentiation and led to a general skeletal hypoplasia without affecting vital organ systems (Almubarak et al., 2021). The rib cage and vertebral column was more affected in these compound homozygous mice compared to the bones in the limb. Given that *Foxc1* and *Foxc2* are abundantly expressed in the condensing limb bud mesenchyme(Almubarak et al., 2021; Hiemisch et al., 1998), such phenotypic differences between the axial and appendicular skeleton were surprising. The mesenchyme that forms the axial skeleton is derived from the sclerotome.

Since the *Col2-cre* transgene is active in sclerotome cells prior to the onset of chondrogenesis but only becomes active in the limb once condensations form (Ovchinnikov et al., 2000), we thought that this earlier timing of the deletion of *Foxc1* and *Foxc2* in axial elements might explain the phenotypic differences we observe in the axial vs appendicular skeleton. To address these issues, we deleted *Foxc1* and *Foxc2* in the limb at an earlier developmental stage than when *Col2-cre* is active using *Sox9-cre* and in *Prx1-cre* mice.

## Results

### Impaired formation of cartilaginous elements in early deletion of *Foxc1* and *Foxc2* in *Sox9-cre* expressing cells

*Foxc1* and *Foxc2* genes are expressed in condensing mesenchyme cells at the onset of chondrogenesis in the limb skeleton (Almubarak et al., 2021; Hiemisch et al., 1998). We examined chondrocyte differentiation (by Alcian blue staining) in embryos engineered to contain floxed alleles of both these transcription factors plus a *Cre* recombinase that had been knocked in to the 3’ UTR of the *Sox9* locus (*Sox9^ires-Cre^*; Akiyama et al., 2005; Sasman et al., 2012). E12.5 embryos that lacked the *Cre* driver displayed Alcian blue staining of both their paraxial mesoderm-derived and appendicular skeletal structures. In contrast, *Sox9^ires-Cre/+^;Foxc1^Δ/Δ^;Foxc2^Δ/Δ^*littermates displayed a dramatic loss of Alcian blue staining in their paraxial mesoderm, but maintained that in their developing limb buds (Fig S1). *Sox9^ires-Cre/+^;Foxc1^Δ/Δ^;Foxc2^+/Δ^*and *Sox9^ires-Cre/+^;Foxc1^+/Δ^;Foxc2^Δ/Δ^* embryos displayed intermediate levels of Alcian blue staining in their paraxial mesoderm. These results indicate that *Foxc1* and *Foxc2* share overlapping roles in promoting chondrogenesis of the paraxial-derived mesoderm. In addition, it suggests that other factors may work in parallel with *Foxc1* and *Foxc2* to promote the initiation of chondrogenesis in the appendicular skeleton.

*Sox9^ires-Cre/+^;Foxc1^Δ/Δ^;Foxc2^Δ/Δ^* mice embryos were not viable after E12.5, preventing further analysis of skeletal development. Therefore, *Prx1-Cre* was used to delete these genes in limb bud mesenchyme (Logan et al., 2002). *Prx1-cre;Foxc1^Δ/Δ^;Foxc2^Δ/Δ^* embryos died shortly after birth and exhibited smaller limbs and feet with abnormal forelimb positioning that resembled decerebrate posture (Fig. 1A, B; white arrow) and exencephaly (yellow arrow). Expression of both *Foxc1* and *Foxc2* mRNA at E16.5 was observed in the skeletal structures of control littermates, but not detected in limb skeletal structures of *Prx1-cre;Foxc1^Δ/Δ^;Foxc2^Δ/Δ^*embryos (Fig. 1C-F), confirming the successful deletion of both genes.

**Fig 1.**
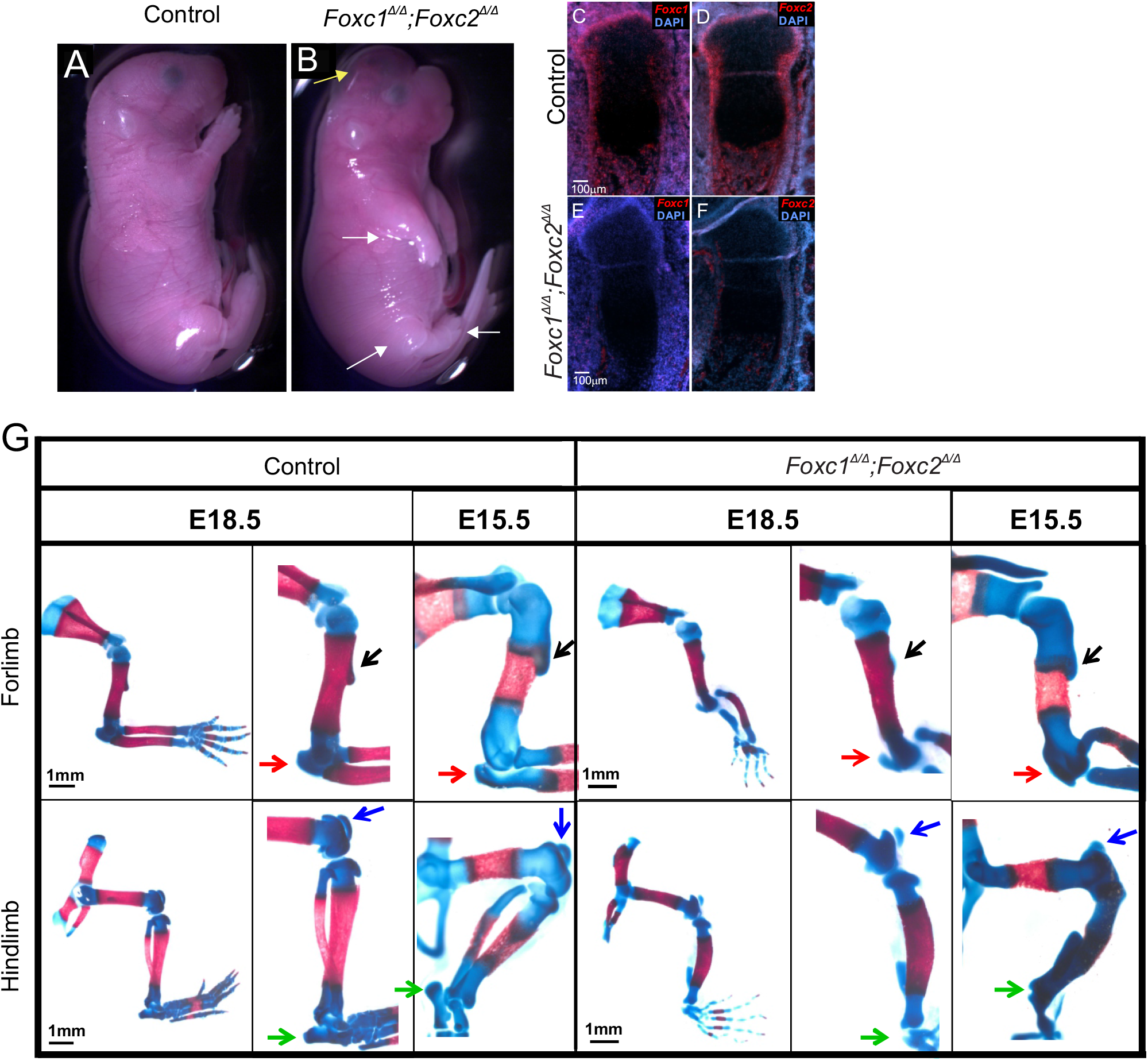
*Foxc1* and *Foxc2* play critical roles in the formation of the distal regions of the appendicular skeleton and support the growth of the bone eminences. (A) E18.5 control and (B) *Prx1-cre;Foxc1^Δ/Δ^;Foxc2^Δ/Δ^* embryos. The *Prx1-cre* mutant mice show the development of exencephaly (yellow arrow) and reduction of forelimbs and hindlimbs size (white arrow).(C-F) In situ hybridization detection of Foxc1 and Foxc2 mRNA expression in control and *Prx1-cre;Foxc1^Δ/Δ^;Foxc2^Δ/Δ^*embryos at E16.5.(G). Whole skeletal staining of the appendicular skeleton in control and *Prx1-cre;Foxc1^Δ/Δ^;Foxc2^Δ/Δ^* embryos at E15.5 and E18.5. Arrows indicate the deltoid tuberosity (black arrow) and olecranon (red arrow) in the forelimb, and the calcaneal tuberosity (green arrow) and the patella (blue arrow) in the hindlimb. Similar results have been obtained with embryos from 9 litters, containing a total of 12 embryos of the least frequently occurring genotype (i.e., *Prx1-Cre; Foxc1^Δ/Δ^;Foxc2^Δ/Δ^*).

Loss of *Foxc1* and *Foxc2* in the developing limb bud drastically affected zeugopod and autopod formation; severely decreasing both the length and thickness of the ulna and fibula in the zeugopod and causing a loss of both wrist and ankle cartilage rudiments and an extreme thinning of the more distal autopod cartilage elements in both the developing fore- and hindlimbs (Fig. 1G). The *Prx1-Cre* mutant limbs showed a severe stunting of skeletal elements in the zeugopod and a thinning or loss of autopod cartilage elements, we also noted that in both the fore- and hindlimbs of *Prx1-cre;Foxc1^Δ/Δ^;Foxc2^Δ/Δ^* embryos, regions of attachment of the skeletal elements to tendons were smaller than in control littermates. In addition, the deltoid tuberosity (black arrow) was initially formed at E15.5 but fails to grow by E18.5. Similarly, the deltoid tuberosity (black arrow), the olecranon (red arrow) in the forelimb; and the calcaneal tuberosity (green arrow) and the patella (blue arrow) in the hindlimb, also fail to significantly grow in *Prx1-cre;Foxc1^Δ/Δ^;Foxc2^Δ/Δ^* embryos (Fig. 1G). These findings signify the importance of *Foxc1* and *Foxc2* in both the proper formation of the zeugopod and autopod cartilage elements and the subsequent development of the bone eminences in the fore- and hindlimbs.

### *Foxc1* and *Foxc2* regulation of endochondral ossification varies temporally during endochondral bone development

Loss of *Foxc1* and *Foxc2* in the developing limb bud affected the distal elements (zeugopod; autopod) more severely than the proximal parts (stylopod). Cartilage formation (Safranin O staining) appeared reduced in the hindlimb autopod of *Prx1-cre;Foxc1^Δ/Δ^;Foxc2^Δ/Δ^* embryos compared to controls at E16.5 (Fig. 2A,B). Next, we investigated the differential proximal-distal phenotypes of the cartilage elements we observed in the mutant mice correlated with a graded expression of *Foxc1* and *Foxc2* along the proximal-distal axis. Expression of *Foxc1* and *Foxc2* mRNA in the hindlimbs at E16.5 was more intense in perichondrium of the autopod compared to that of either the tibia (in the zeugopod) or the femur (in the stylopod; Fig. 2C, D). At E14.5, expression of both *Foxc1* and *Foxc2* mRNA signals were prominent in the perichondrium and the growth plate, including the hypertrophic chondrocytes (Fig. 2G, H). Like the E16.5 limbs, the E14.5 limbs also displayed increased *Foxc1* and *Foxc2* in the autopod compared to the proximal bone elements. Thus the expression of Foxc1 and Foxc2 is most intense in the newly formed perichondrium of more distal skeletal elements; but is also expressed in the growth plate of the POC.

**Fig 2.**
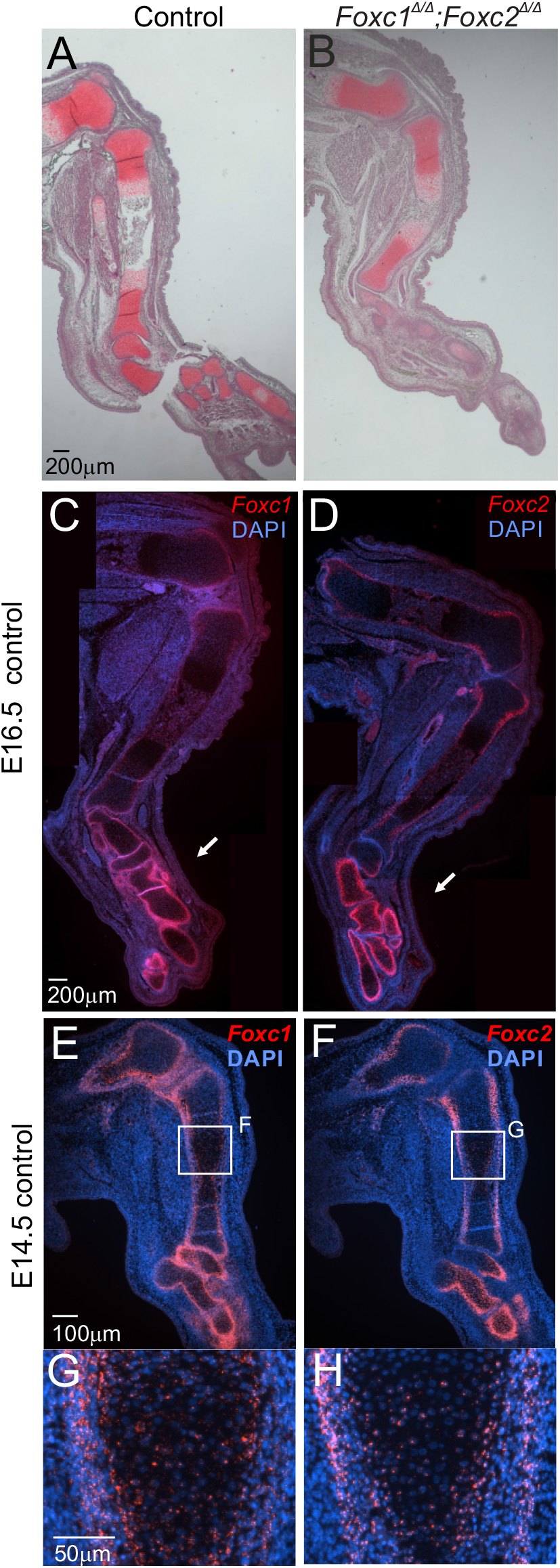
Elevated expression of *Foxc1* and *Foxc2* mRNAs in distal forelimb structures. Delayed cartilage formation in hindlimb autopod structures. Safranin O staining of Control (A) and *Prx1-cre;Foxc1^Δ/Δ^;Foxc2^Δ/Δ^* (B) hindlimb at E16.5. Expression of *Foxc1* and *Foxc2* mRNAs hindlimb sections at E16.5 (C,D) and E14.5 (E-H).

### Loss of *Foxc1* and *Foxc2* reduced proliferation during early growth plate development

We tested whether reduced chondrocyte proliferation accounted for the reduced size of the mutant limbs. Proliferation of cells in both control and *Prx1-cre;Foxc1^Δ/Δ^;Foxc2^Δ/Δ^* growth plates was determined by KI67 IF, which labels actively dividing cells (Gerdes et al., 1984). Analysis of control limbs revealed the highest number of KI67-positive cells in the proximal growth plate of the tibia at E14.5 in comparison to later time points (Fig. 3A-C). This finding indicates that proliferative chondrocytes were more abundant at E14.5 than later time points in the tibia cartilage element. Interestingly, the growth plate of the tibia in *Prx1-cre;Foxc1^Δ/Δ^;Foxc2^Δ/Δ^* E14.5 exhibited an approximate 50% reduction in the number of KI67-positive chondrocytes compared to the control limbs (Fig. 3C). However, no changes in proliferation activity were detected in control and mutant limbs at E15.5 and E16.5. This result suggests that *Foxc1* and *Foxc2* act at a narrow window (prior to E15.5) to regulate chondrocyte proliferation.

**Fig 3.**
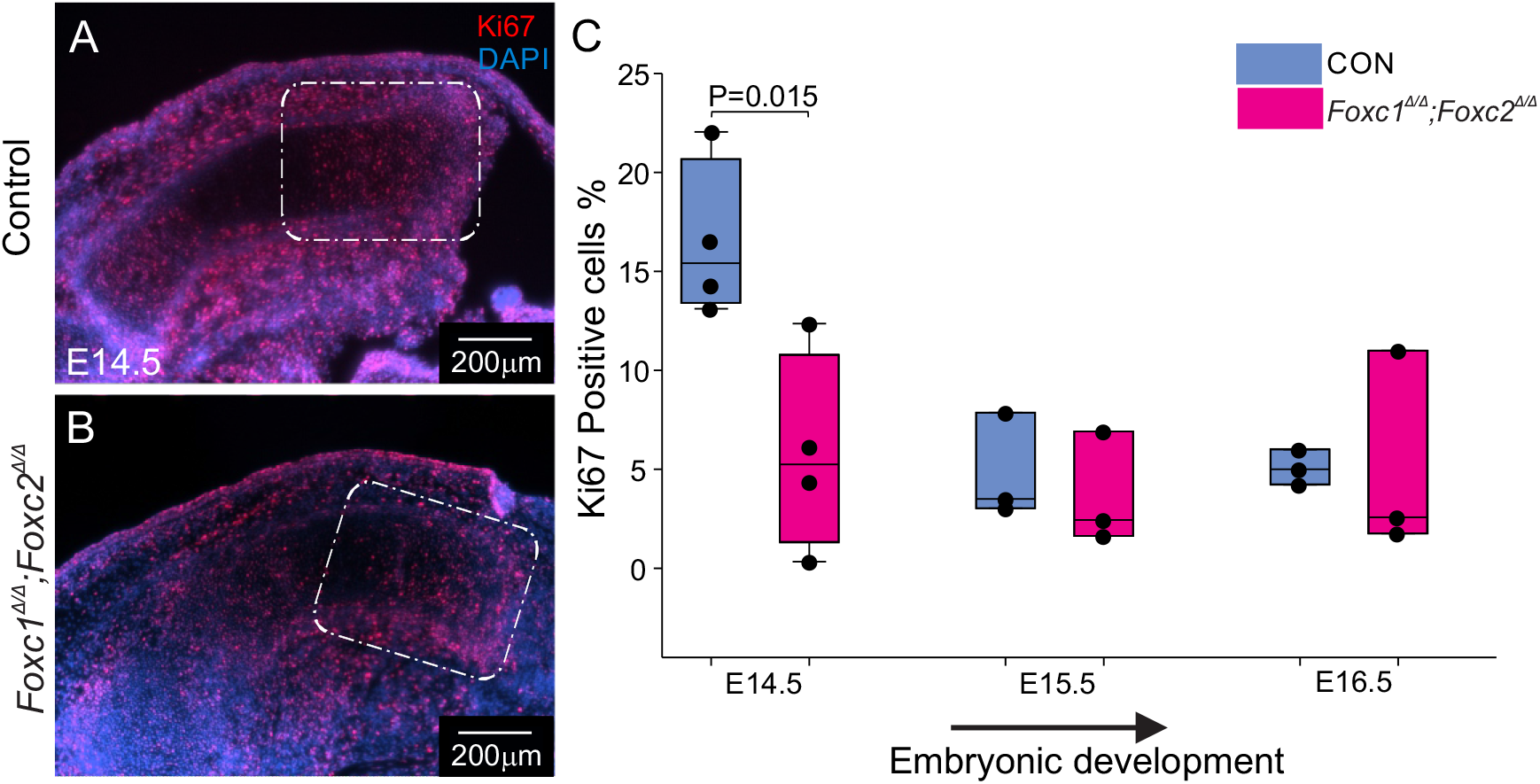
Chondrocyte proliferation is reduced in *Prx1-cre;Foxc1^Δ/Δ^;Foxc2^Δ/Δ^* limbs at E14.5 but not affected at later stages. Cell proliferation in the proximal tibia growth plate was assessed by KI67 IF. Representative micrographs for control (A) and *Prx1-cre;Foxc1^Δ/Δ^;Foxc2^Δ/Δ^* (B) proximal tibia sections are shown (white box). (C) The percentage of KI67-positive cells was determined in control and *Prx1-cre;Foxc1^Δ/Δ^;Foxc2^Δ/Δ^* proximal tibia growth plates at E14.5, E15.5 and E16.5 tibia. Data presented are from 3-4 embryos per genotype.

To investigate why proliferation was reduced in *Prx1-cre;Foxc1^Δ/Δ^;Foxc2^Δ/Δ^* chondrocytes, we examined the expression of IHH and PTHRP signaling components that regulate chondrocyte proliferation in the growth plate at E14.5. We detected prominent expression of *Ihh* in the prehypertrophic chondrocytes and the newly forming hypertrophic chondrocytes in the developing tibia of control littermates E14.5 (Fig. 4A). In contrast, the size of the *Ihh* expressing region in the developing tibia of *Prx1-cre;Foxc1^Δ/Δ^;Foxc2^Δ/Δ^* embryos was greatly reduced (Figs. 4B and Q). *Ptch1* and *Ptch2* are receptors for IHH and their expression is elevated in response to ligand binding; thus their expression can be used to monitor active hedgehog signalling (Alman, 2015). *Ptch1* mRNA was detected in both the proliferating zone (PZ) chondrocytes and in the perichondrium that surrounded *Ihh-*expressing cells in the tibia of E14.5 control embryos (Fig 4D). In the *Prx1-cre;Foxc1^Δ/Δ^;Foxc2^Δ/Δ^* embryos, *Ptch1* expression appeared reduced but was localized in a similar pattern, that surrounded the smaller *Ihh-*expressing region (Fig 4E). *Ptch2* expression was predominant in the perichondrium in the control tibias at E14.5, while expression in *Prx1-cre;Foxc1^Δ/Δ^;Foxc2^Δ/Δ^*embryos was less intense (Fig 4F-G). In the control embryos, *Gli1, Gli2, Gli3* mRNAs were detected in the resting zone and proliferating zone chondrocytes as well as the perichondrium. While localization of these transcripts was similar in the *Prx1- cre;Foxc1^Δ/Δ^;Foxc2^Δ/Δ^* mutants, the intensity of their expression was lower, correlating with decreased expression of *Ihh* (Fig 4H-M). *Pthlh* displayed comparable expression in control and *Prx1-cre;Foxc1^Δ/Δ^;Foxc2^Δ/Δ^* embryos, with transcripts localizing to the resting zone chondrocytes (Fig. 4N, O). The Parathyroid hormone receptor (*Pth1r*) was expressed in a smaller domain within the tibial growth plate of *Prx1-cre;Foxc1^Δ/Δ^;Foxc2^Δ/Δ^*embryos compared to that in a control littermate (Fig. 4, P, Q, and R). Collectively, these data indicate that the absence of *Foxc1* and *Foxc2* slows the formation of *Ihh-*expressing pre-hypertrophic chondrocytes, but that the signaling components needed to mediate IHH-PTHLH signalling are functioning in *Prx1- cre;Foxc1^Δ/Δ^;Foxc2^Δ/Δ^* embryos at E14.5.

**Fig 4.**
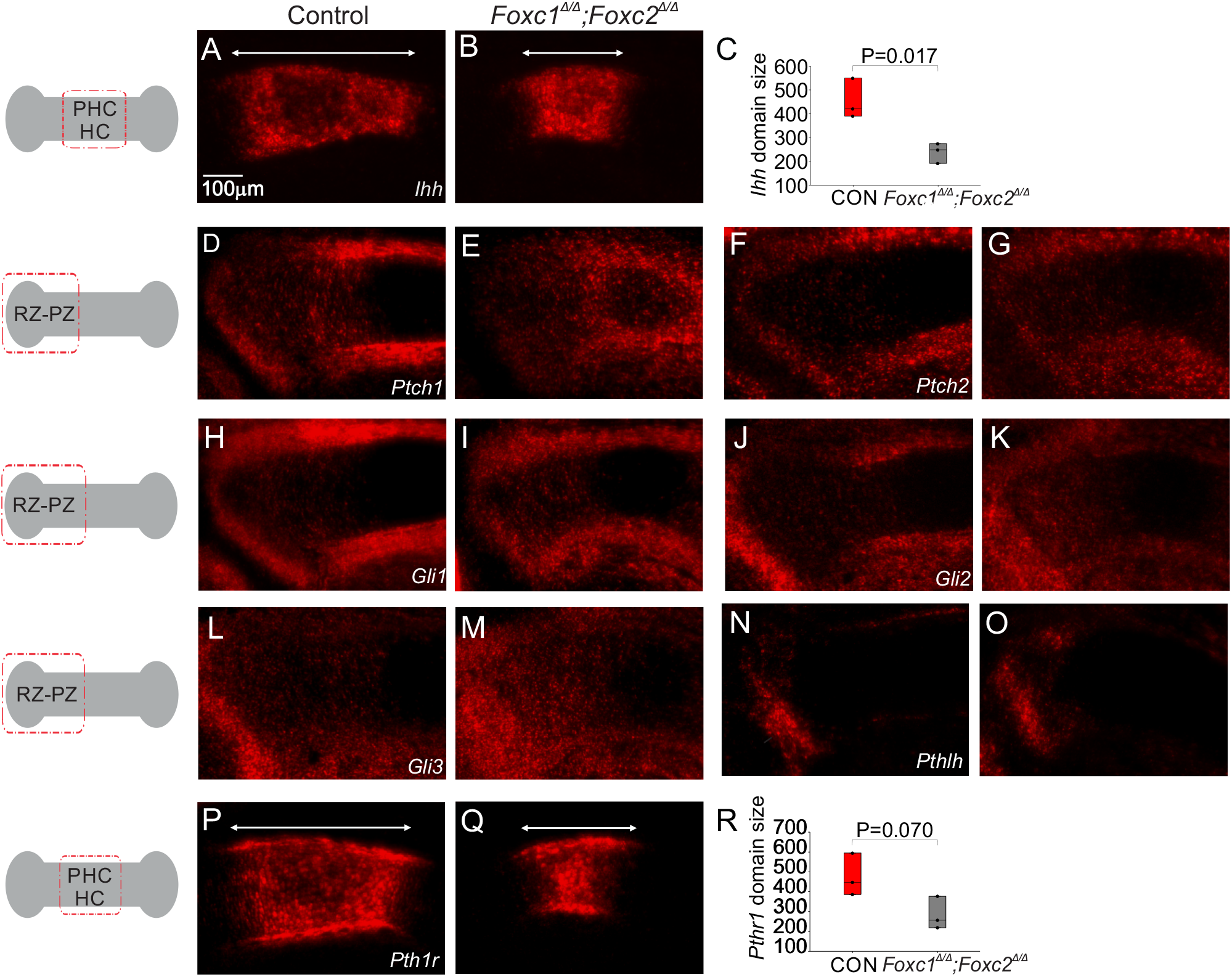
*Ihh* expression expression is reduced in *Prx1-cre;Foxc1^Δ/Δ^;Foxc2^Δ/Δ^* prehypertrophic chondrocytes at E14.5. Assessment of the IHH-PTHLH signalling axis function in *Prx1-cre;Foxc1^Δ/Δ^;Foxc2^Δ/Δ^* embryos. Expression of *Ihh* (A-C), *Ptch1* (D,E), *Ptch2* (F,G), *Gli2*(H,I), *Gli1*(J,K), *Gli3* (L,M), *Pthlh* (N,O), *Pth1r* (P-R) was assessed by in situ hybridization in the tibia of control and *Prx1- cre;Foxc1^Δ/Δ^;Foxc2^Δ/Δ^*embryos at E14.5. The position in the growth plate for micrographs is indicated in the schematic diagram on the left. Scale bar 100 μm. PHC-prehypertrophic chondrocytes; HC-hypertrophic chondrocytes; RZ-resting zone; PZ-proliferating zone. Statistical analysis performed using Student’s t-test from three littermate pairs (control and mutant).

### Expanded hypertrophic chondrocyte zone in *Prx1-cre;Foxc1^Δ/Δ^;Foxc2^Δ/Δ^* embryos

To evaluate whether the delayed maturation of prehypertrophic chondrocytes in limbs lacking Foxc1 and Foxc2 persisted later in development, we evaluated chondrogenic differentiation and growth plate organization in *Prx1-cre;Foxc1^Δ/Δ^;Foxc2^Δ/Δ^* embryos and in control littermates at later stages of development. In contrast to the tibial growth plates of E14.5 *Prx1-cre;Foxc1^Δ/Δ^;Foxc2^Δ/Δ^*embryos, which displayed delayed formation of *Ihh-*expressing pre- hypertrophic chondrocyte, at E15.5 we did not observe any anomalies in the organization of the growth plate in this same cartilage element (Fig. S2). SOX9 and SOX6 protein localization in the resting zone (RZ) and proliferating zone (PZ) was similar between the control and *Prx1- cre;Foxc1^Δ/Δ^;Foxc2^Δ/Δ^* growth plates at E15.5 (Fig. S2A-D); and localization of *Fgfr3* RNA confirmed that formation of the PZ chondrocytes formed in E15.5 *Prx1-cre;Foxc1^Δ/Δ^;Foxc2^Δ/Δ^*embryos (Fig. S2E, F). Finally, expression of *Ihh* mRNA was detected at comparable levels in the prehypertrophic chondrocytes in the proximal tibial growth plate of E15.5 *Prx1- cre;Foxc1^Δ/Δ^;Foxc2^Δ/Δ^* embryos (Fig. S2G, H). Consistent with the relatively normal tibial growth plates we observed in E15.5 *Prx1-cre;Foxc1^Δ/Δ^;Foxc2^Δ/Δ^* embryos, no anomalies were detected in the formation of the RZ, PZ and prehypertrophic chondrocytes (PHC) of the tibia growth plates in E16.5 mutant embryos (Fig 5). We found that *Fgfr1* and *Fgfr3 mRNAs* were detected in similar patterns in the RZ and PZ chondrocytes, respectively, in both E16.5 control and mutant growth plates (Fig. 5A-D). In addition, both *Ihh* mRNA and RUNX2 protein were similarly localized to the pre-hypertrophic chondrocyte zone in both control and mutant limbs at E16.5 (Fig. 5E-H). However, some RUNX2 signal was detected in the hypertrophic chondrocytes in the E16.5 *Prx1-cre;Foxc1^Δ/Δ^;Foxc2^Δ/Δ^* growth plate but not in the controls (Fig. 5E, F). Interestingly, at E16.5 we observed an expanded COLX IF signal in the hypertrophic chondrocyte zone in the absence of *Foxc1* and *Foxc2* (Fig. 5I, J). However this expansion was not a result of increased *Colx* expression, as *m*RNA levels were similar in both the control and the *Prx1-cre;Foxc1^Δ/Δ^;Foxc2^Δ/Δ^*hypertrophic zone (Fig. 5K, L). Taken together, our findings suggest that the initial delayed maturation of PHC in limbs lacking *Foxc1* and *Foxc2* (at E14.5) is normalized at later stages of development; and that COLX protein was persistent in growth plates of *Prx1-cre;Foxc1^Δ/Δ^;Foxc2^Δ/Δ^* embryos, likely from impaired hypertrophic chondrocyte turnover.

**Fig 5.**
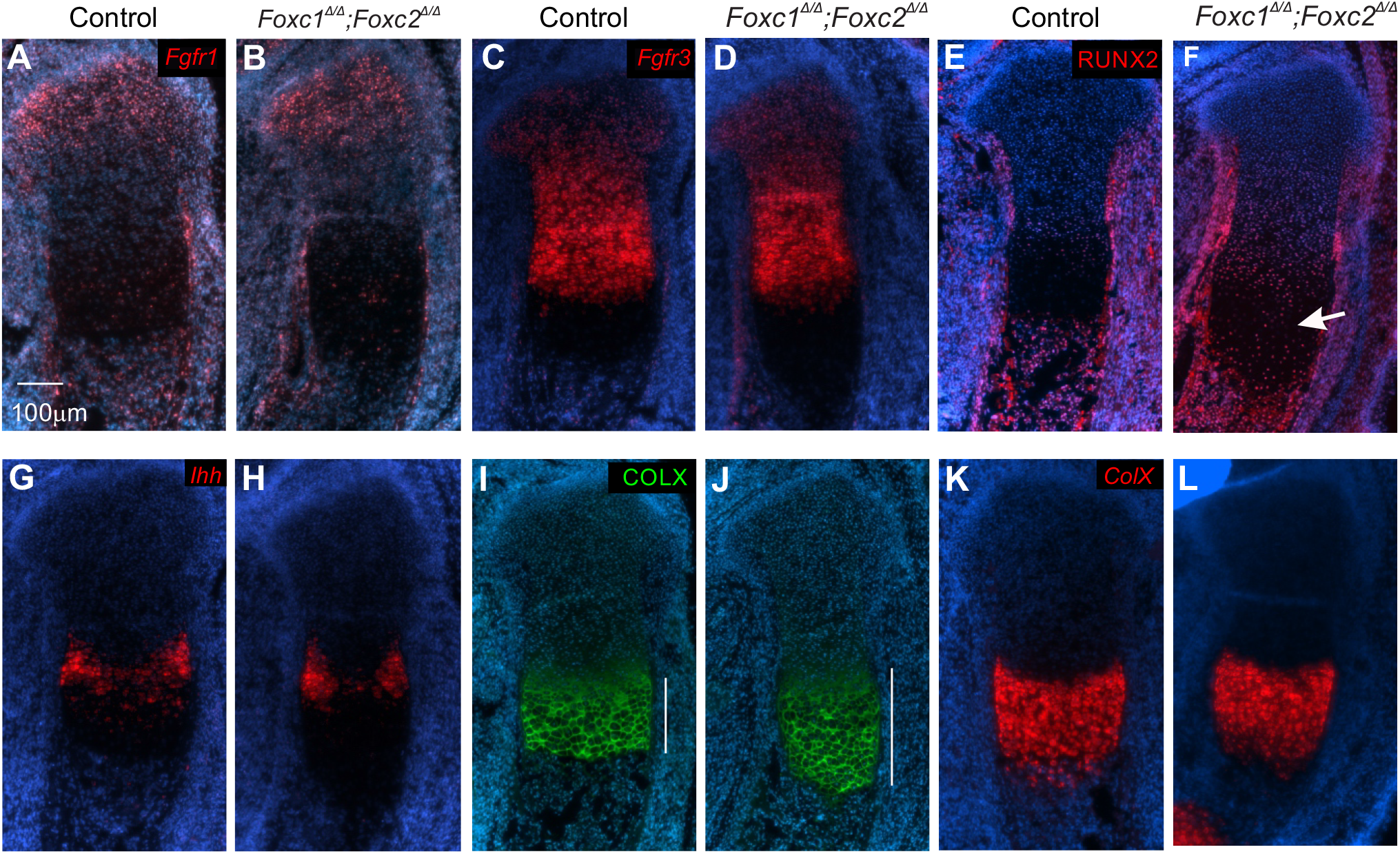
Deletion *Foxc1* and *Foxc2* disrupted the terminal hypertrophic chondrocyte differentiation. Chondrocyte differentiation and growth organization was examined in proximal tibias at E16.5 from control and using *Prx1-cre;Foxc1^Δ/Δ^;Foxc2^Δ/Δ^* embryos. (A,B) *Fgfr1* (resting zone), (C,D) *Fgfr3* (columnar chondrocytes), (E,F) RUNX2 (pre-hypertrophic chondrocytes), (I,J) COLX protein (hypertrophic chondrocytes), and (K,L) *Colx* mRNA

### Entry and exit from chondrocyte hypertrophy is delayed in *Prx1-cre;Foxc1^Δ/Δ^;Foxc2^Δ/Δ^* mice

The reduction of the *Ihh-*expressing pre-hypertrophic chondrocyte zone at E14.5, and the expansion of the hypertrophic chondrocytes zone at E16.5 point to a primary defect in hypertrophic chondrocyte differentiation. Thus, we decided to track the chondrogenic differentiation progression in the growth plate from E14.5-E17.5 by monitoring COL2 and COLX protein localization. We observed no differences in the length of COL2-expressing regions between the control and mutant limb sections at all time points examined (Fig. S3A-P). Moreover, the distance between the RZ and HCs were not affected in the *Prx1- cre;Foxc1^Δ/Δ^;Foxc2^Δ/Δ^*embryos, indicating that the IHH-PTHLH singling network was functioning. The size of the COLX-expression domain was reduced in the hypertrophic chondrocyte zone at E14.5 and E15.5 in *Prx1-cre;Foxc1^Δ/Δ^;Foxc2^Δ/Δ^* tibias (Fig. 6A-F). At later ages we observed an expansion of the proximal and distal COLX-expressing hypertrophic chondrocyte domains at E16.5 and E17.5 compared to the control limbs (Fig. 6G-O). In addition, the primary ossification center (POC), located between the two extended hypertrophic chondrocyte domains, at E16.5 and E17.5 was much smaller in *Prx1-cre;Foxc1^Δ/Δ^;Foxc2^Δ/Δ^*limbs compared to the widely formed POC in control limbs (Fig. 6G-P). These data suggest that the absence of *Foxc1* and *Foxc2* in chondrocytes caused both a delayed entry into hypertrophy followed by a delayed exit; thereby leading to the formation of a smaller primary ossification center.

**Fig 6.**
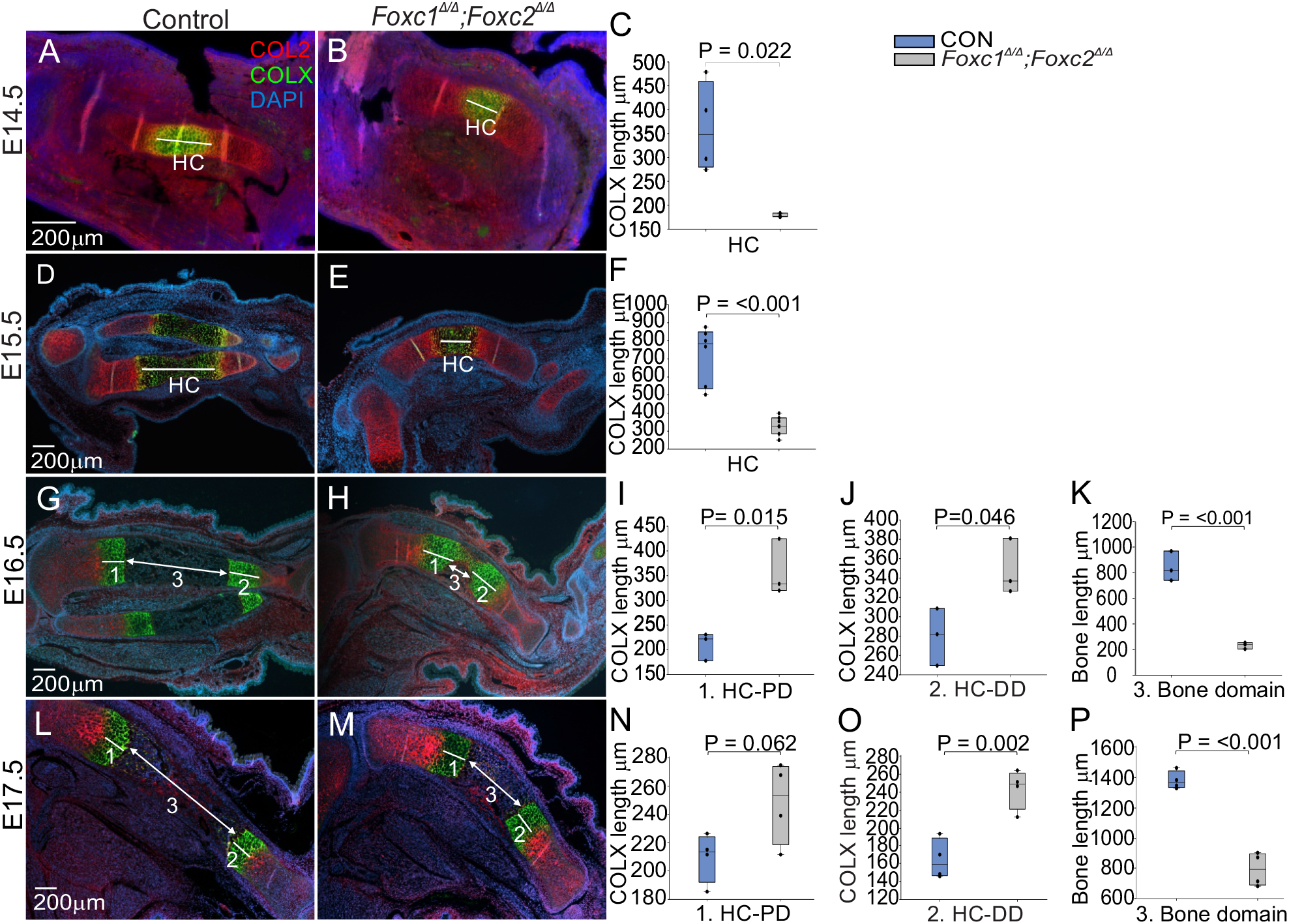
Delayed entry into and exit from chondrocyte hypertrophy in *Prx1- cre;Foxc1^Δ/Δ^;Foxc2^Δ/Δ^* embryos. Formation and growth of the hypertrophic chondrocyte zone was analyzed by measuring the length of COLX-positive cells (green) at E14.5 (A-C), E15.5 (D-F), E16.5 (G-K) and E17.5 (L- P). The size of the primary ossification center was determined by measuring the distance between terminal COLX expression zones at E16.5 and E17.5. 1-Hypertrophic chondrocytes proximal domain (1-HC-PD), 2- COLX-positive Hypertrophic chondrocytes distal domain (2- HC-DD). 3- Bone domain/primary ossification center. Scale bar 200 μm. Statistical analysis was performed using Student t-test. (n=4).

To test whether the expansion of the hypertrophic chondrocytes zone at E16.5 in *Prx1- cre;Foxc1^Δ/Δ^;Foxc2^Δ/Δ^* mice was due to a reduction in cell death, we performed terminal deoxynucleotidyl transferase dUTP nick end labeling (TUNEL) *in situ hybridization* (TUNEL) at E15.5 and E16.5. At E15.5, apoptosis was detected in a population of cells adjacent to the perichondrium bordering the hypertrophic chondrocytes in both control and mutant limbs (white arrow) (Fig. 4S A, B). Cell death was not detected within the growth plate chondrocytes (Fig. 4S A, B, yellow asterisk). At a later stage, TUNEL positive cells were found among the perichondrium and primary POC (Fig. 4S C, green asterisk). However, in *Prx1- cre;Foxc1^Δ/Δ^;Foxc2^Δ/Δ^* mutant limbs, cell death was primarily limited to the perichondrium and detected in only a small region of the nascent POC (Fig. 4S D, green asterisk). Since very few TUNEL-positive cells were detected in the HC in both the control and mutant tibia suggests that the changes in the length of the HC zone we observed in the *Prx1-cre;Foxc1^Δ/Δ^;Foxc2^Δ/Δ^* embryos was not a consequence of impaired cell death in this region.

### *Foxc1* and *Foxc2* are required for proper mineralization and bone formation

We next examined if the absence of *Foxc1* and *Foxc2* has affected osteoblast formation and bone mineralization. In the tibia of E15.5 *Prx1-cre;Foxc1^Δ/Δ^;Foxc2^Δ/Δ^* embryos, the COL1 signal was reduced in the osteochondral junction compared to control littermates; and MMP13 was absent in the HC and the emerging POC of the mutant embryos (Fig 7A,D). OSX was detected in the bone collar and the PHC in the *Prx1-cre;Foxc1^Δ/Δ^;Foxc2^Δ/Δ^* tibia sections, but was not detected in the future POC (Fig. 7 B, E). In addition, mineralization was also impaired at this stage, as we observed no detectable Von Kossa staining in the tibia of E15.5 *Prx1- cre;Foxc1^Δ/Δ^;Foxc2^Δ/Δ^*embryos compared to the control embryos (Fig. 7 C,F). The impaired osteoblast formation and mineralization defect persisted at E16.5. COL1 localization was detected in the osteochondral junction, the bone collar and POC in control tibias, while in mutants, COL1 was detected in the osteochondral junction and in a group of cells in the nascent POC, connecting the proximal and distal HC (Fig. 7 G,J). MMP13 was restricted to the osteochondral junction of the control tibias, while a broader protein distribution was detected in the HC and POC of *Prx1-cre;Foxc1^Δ/Δ^;Foxc2^Δ/Δ^* embryos (Fig. 7 G,J). OSX was detected in the bone collar and POC (Fig. 7H, K), however the POC was noticeably smaller in *Prx1- cre;Foxc1^Δ/Δ^;Foxc2^Δ/Δ^*embryos. In addition, Von Kossa staining revealed persistent mineralization of hypertrophic chondrocytes and an asymmetrical staining localize to the posterior bone collar (black arrow), as well as reduced mineralization in the POC in the *Prx1- cre;Foxc1^Δ/Δ^;Foxc2^Δ/Δ^* embryos compared to controls (Fig. 7 I, L). At E17.5, osteoblast formation and mineralization continued to be reduced in *Prx1-cre;Foxc1^Δ/Δ^;Foxc2^Δ/Δ^* limbs. MMP13, COL1 (Fig. 7 M,P) and OSX (Fig. 7 N, Q) were localized to a smaller POC in *Prx1- cre;Foxc1^Δ/Δ^;Foxc2^Δ/Δ^* embryos. Collectively, these data revealed that osteoblast formation and mineralization of the bone still occurred in the absence of *Foxc1* and *Foxc2* in limb bud mesenchyme, but that these transcription factors are essential for proper mineralization and formation of osteoblasts in the trabecular bone.

**Fig. 7.**
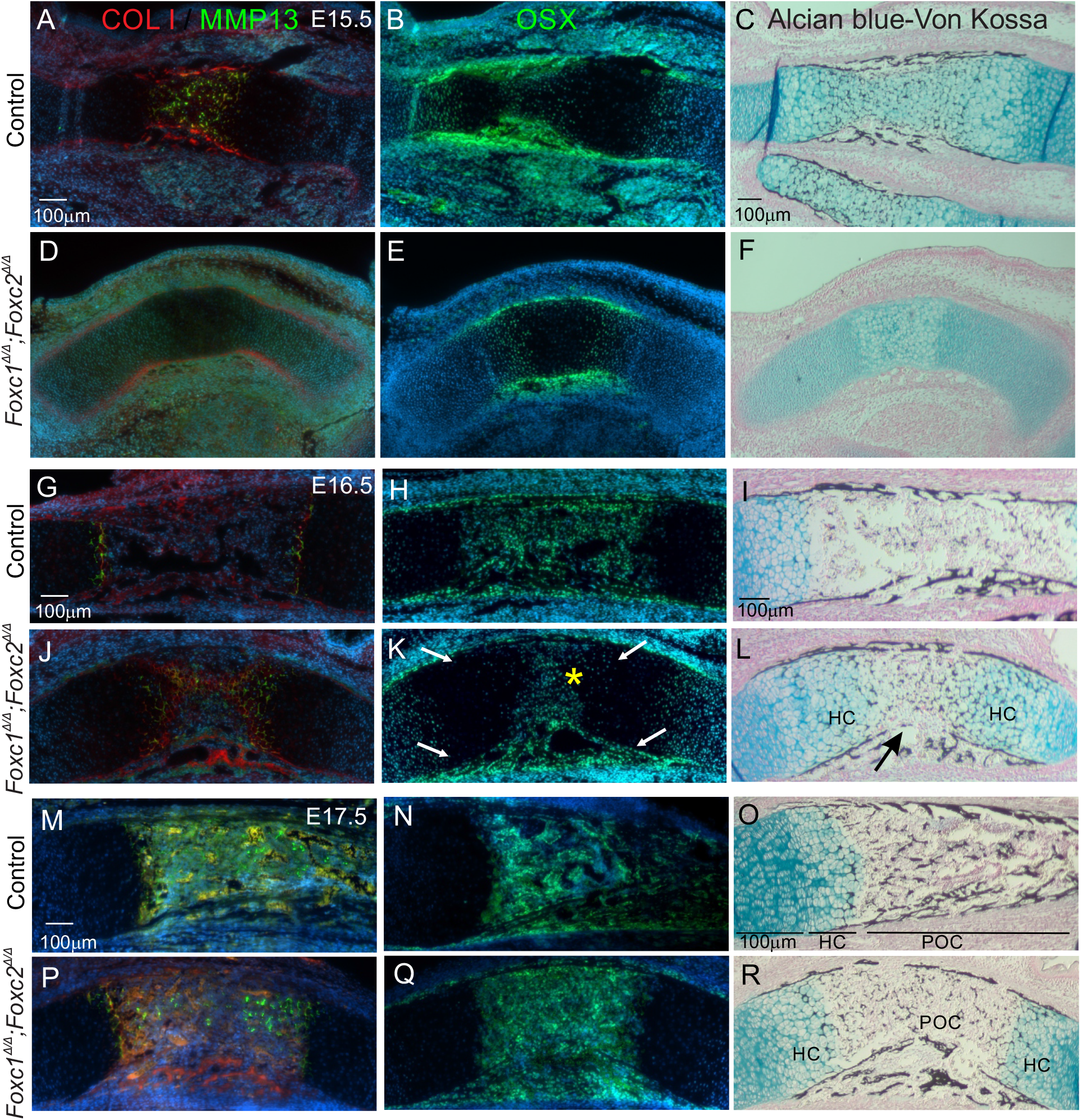
Delayed osteoblast differentiation and mineralization in the POC of *Prx1- cre;Foxc1^Δ/Δ^;Foxc2^Δ/Δ^* embryos. Osteoblast formation, hypertrophic chondrocyte remodelling and mineralization was examined at E15.5 (A-F), E16.5 (G-L), and E17.5 (M-R) by immunofluorescent detection of COL1, OSX (osteoblast differentiation), and MMP13 (hypertrophic chondrocytes in the primary ossification center of control and *Prx1- cre;Foxc1^Δ/Δ^;Foxc2^Δ/Δ^* mice. Mineralization was detected through Von Kossa staining . Arrows denote asymmetrical ossification from posterior side of the bone collar in *Prx1- cre;Foxc1^Δ/Δ^;Foxc2^Δ/Δ^* embryos. HC-hypertrophic chondrocytes; POC-primary ossification center. Scale bar 100 μm (N=3)

### Bone remodeling was compromised by the absence of *Foxc1* and *Foxc2* in long bones

Since the formation of the POC was disrupted in *Prx1-cre;Foxc1^Δ/Δ^;Foxc2^Δ/Δ^* mutants, yet osteoblast differentiation and mineralization occurred in the periosteum/perichondrium, we examined whether impaired bone remodeling or vascularization prevented invasion of osteoblasts into the bone interior of the *Prx1-cre;Foxc1^Δ/Δ^;Foxc2^Δ/Δ^* limbs. We visualized osteoclasts using Tartrate-Resistant Acid Phosphatase (TRAP) staining at E16.5 and E17.5 in the control and *Prx1-cre;Foxc1^Δ/Δ^;Foxc2^Δ/Δ^* limbs. TRAP-positive cells were localized in the perichondrium, periosteum, and the osteochondral junction in the control tibia POC. However, there were fewer osteoclasts in the *Prx1-Cre* mutants at both time points (Fig. 8 A-E). The low number of osteoclasts in the mutants may cause slower bone resorption activity leading to reduced bone remodeling and formation of shorter limbs (Lademann et al., 2020). We then monitored expression of *Tnfsf11* mRNA (RANKL) which stimulates osteoclast recruitment.

**Fig. 8.**
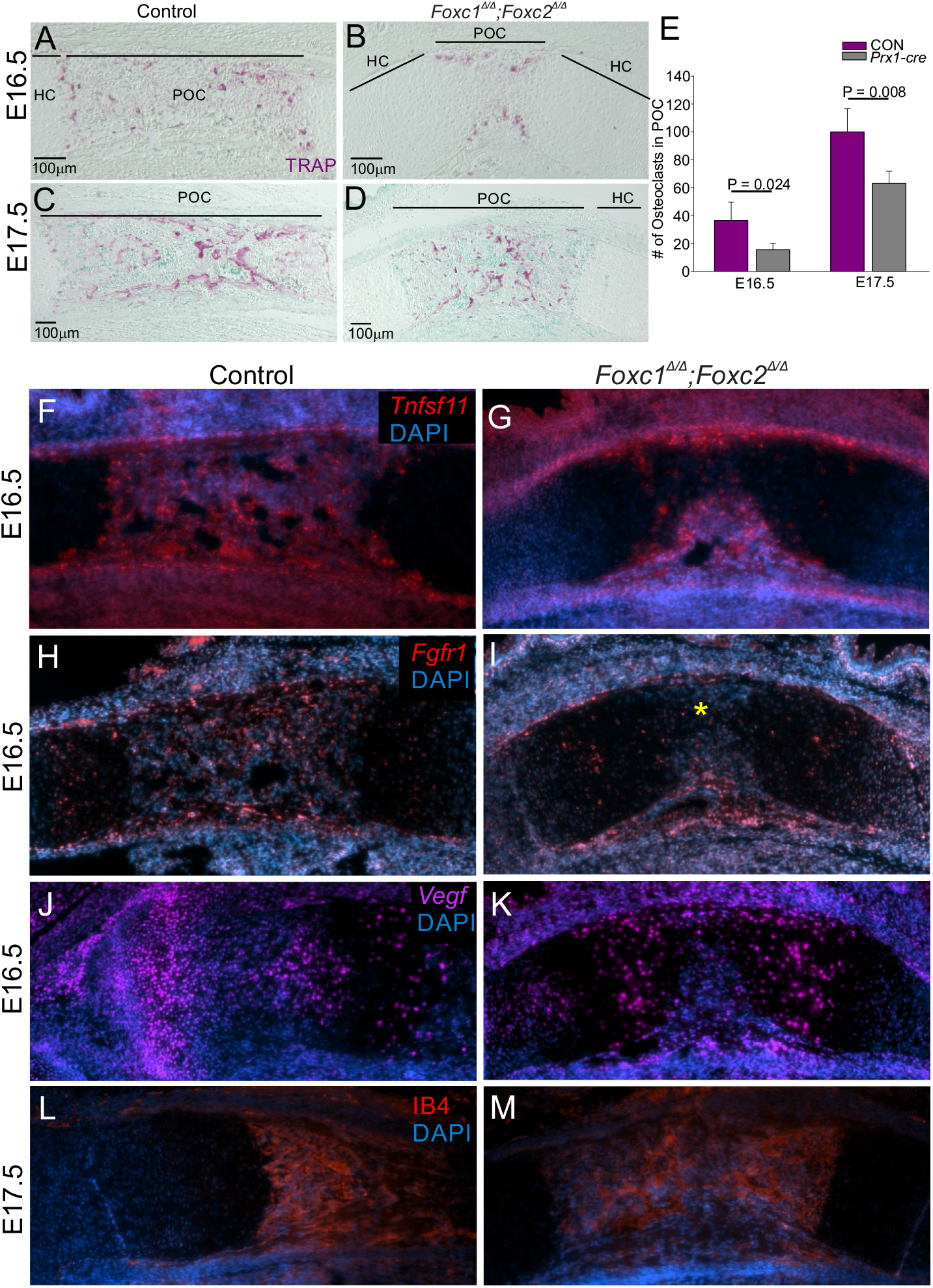
Reduced activation of osteoclasts in the POC of *Prx1-cre;Foxc1^Δ/Δ^;Foxc2^Δ/Δ^* embryos. TRAP staining was performed on E16.5 (A, B) and E17.5 (C, D) control and *Prx1- cre;Foxc1^Δ/Δ^;Foxc2^Δ/Δ^* tibia to assess osteoclast localization in the primary ossification center. Widespread TRAP signal was distributed throughout the POC of control embryos (A). In contrast, TRAP signal was localized at the bone collar in mutants (B). At E17.5, osteoclasts had a comparable distribution pattern throughout the POC in both control and mutant limbs (C, D). Fewer TRAP-positive cells were detected in the *Prx1-cre;Foxc1^Δ/Δ^;Foxc2^Δ/Δ^* limb at E16.5 and E17.5 in comparison to their controls (E). Expression of *Tnfsf11* (RANKL) mRNA was detected in the control and *Prx1-cre;Foxc1^Δ/Δ^;Foxc2^Δ/Δ^* POC (F, G). *Fgfr1* mRNA expression was reduced in the POC of *Prx1-cre;Foxc1^Δ/Δ^;Foxc2^Δ/Δ^* embryos (yellow asterisk; H, I). *Vegfa* mRNA was localized equally in the HC and POC in control and *Prx1-cre;Foxc1^Δ/Δ^;Foxc2^Δ/Δ^*limbs (J, K). IB4 was detected in both E17.5 control and *Prx1-cre;Foxc1^Δ/Δ^;Foxc2^Δ/Δ^*POC (L, M). HC- hypertrophic chondrocytes; POC-primary ossification center. Statistical analysis was performed by the Student’s t-test. Error bars indicate standard deviation. (n=3).

Expression of *Tnfsf11* was detected in osteoblasts in control and *Prx1-cre;Foxc1^Δ/Δ^;Foxc2^Δ/Δ^* mutant limbs, although the expression region was much smaller in the mutant limbs, likely owing to the reduction of osteoblast formation (Fig. 8 F, G). Other regulators of osteoclast formation, such as *Fgfr1(Lu et al., 2009)*was also reduced in the POC of the *Prx1- cre;Foxc1^Δ/Δ^;Foxc2^Δ/Δ^* embryos (Fig. 8 H, I).

Angiogenesis and blood vessel invasion into the POC are necessary for bone formation (Sivaraj and Adams, 2016). Thus, we assessed whether vascularization was affected in the limbs of *Prx1-cre;Foxc1^Δ/Δ^;Foxc2^Δ/Δ^*embryos. *Vegfa* mRNA was localized in the hypertrophic chondrocytes in both control and the *Prx1-cre* mutant limbs, with more *Vegfa* signal marking the extended hypertrophic chondrocytes zone in the *Prx1-cre;Foxc1^Δ/Δ^;Foxc2^Δ/Δ^* E16.5 tibia (Fig. 8 J, K). Moreover, vascular endothelial marker isolectin-B4 (IB4) localization confirmed the presence of blood vessels in both control and mutant POCs (Fig. 8 L, M). Collectively, these results suggest that expression of *Foxc* transcription factors in limb bud mesenchyme play an essential role in facilitating osteoclast recruitment and activation, which is necessary for proper bone formation and resorption. In contrast, deletion of *Foxc1* and *Foxc2* in limb bud mesenchymal cells did not block blood vessel invasion into the bone cavity.

### *Foxc1* is required for *Phex* expression to maintain bone mineralization

We detected OPN protein in the bone collar and the primary ossification center in both control and *Prx1-cre;Foxc1^Δ/Δ^;Foxc2^Δ/Δ^*limbs (Fig. 9A-D). However, there was a more intense OPN signal located within the small POC in the mutant limbs at E16.5 and E17.5 (Fig. 9B, D). OPN is a glycoprotein that is expressed in hypertrophic chondrocytes, osteoblasts, and osteoclasts. In order for mineralization to proceed, OPN is proteolytically processed and degraded by enzymes such as Phosphate Regulating Endopeptidase Homolog X-linked (PHEX; (Addison et al., 2010; Zurick et al., 2013). We previously found that *Phex* mRNA levels were reduced in Col2-cre*;Foxc1^Δ/Δ^;Foxc2^Δ/Δ^* mice, as determined by RNA-seq analysis. Consistent with this prior finding, we similarly found a dramatic reduction in *Phex* expression in the POC of *Prx1-cre;Foxc1^Δ/Δ^;Foxc2^Δ/Δ^* tibias at E16.5 and E17.5 compared to controls (Fig. 9E-H). In addition, areas where Phex expression was reduced in *Prx1-cre;Foxc1^Δ/Δ^;Foxc2^Δ/Δ^*limbs overlapped with areas where OPN levels were elevated. Notably, the expression of both *Foxc1* and *Phex* mRNA significantly overlaps in the POC at E16.5 and E17.5 in control limbs (Fig. 9M-P). These results indicate that *Foxc1* and *Foxc2* are required for the expression of *Phex* in the POC. The consequent absence of *Phex* expression may lead to improper processing of OPN; and thus block the proper mineralization in the POC of *Prx1-cre;Foxc1^Δ/Δ^;Foxc2^Δ/Δ^*mutants.

**Fig. 9.**
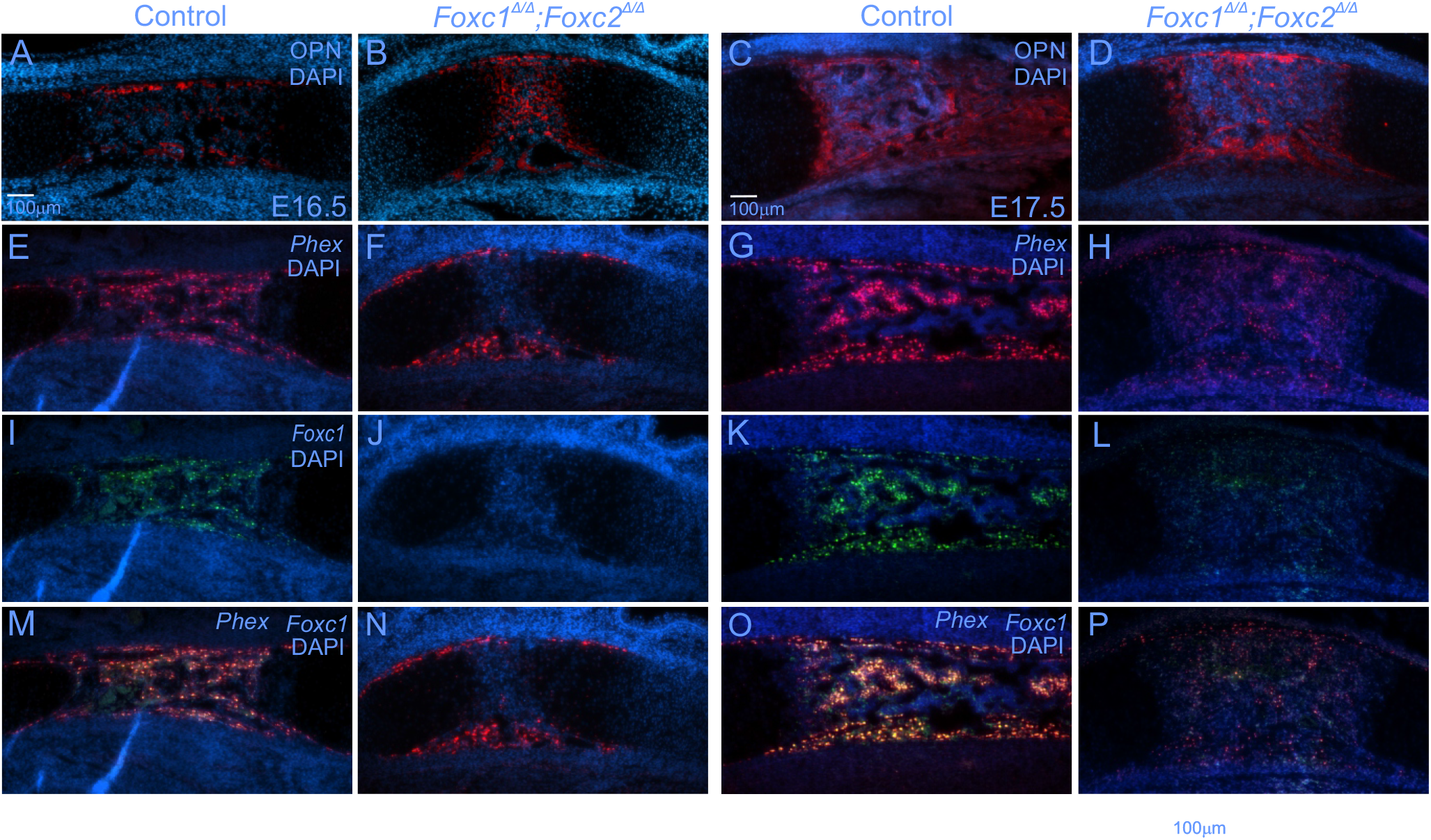
Foxc1 and Foxc2 are required for *Phex* expression in the POC. OPN levels were determined in the POC of the control and *Prx1-cre;Foxc1^Δ/Δ^;Foxc2^Δ/Δ^*tibia at E16.5 (A,B) and E17.5 (C,D). *Phex* mRNA expression was determined by *in situ* hybridization at E16.5 (E,F) and E17.5 (G,H) in control and *Prx1-cre;Foxc1^Δ/Δ^;Foxc2^Δ/Δ^*tibia. Comparison of *Phex* mRNA localization with *Foxc1* mRNA expression was determined by in situ hybridization at E16.5 (I,JM,N) and E17.5 (K,L,O,P)) in the control and mutant tibia.

## Discussion

Foxc1 and Foxc2 are expressed in the condensing mesenchyme that form skeletal structures, suggesting that these transcription factors may participate in the early stages of bone formation (Almubarak et al., 2021; Hiemisch et al., 1998). To understand how *Foxc1* and *Foxc2* control early steps of endochondral ossification, we generated two conditional compound knockouts using *Sox9- cre* or *Prx1-cre* to delete *Foxc1* and *Foxc2* in chondrocyte progenitors and limb bud osteochondral progenitors, respectively. Loss of *Foxc1* and *Foxc2* function in the *Sox9* lineage, resulted in the near absence of chondrogenesis in the vertebral column and a reduction in the size of the cartilaginous elements in the limb buds at E12.5. This difference in the effect of Foxc1/Foxc2 loss in Sox9-expressing cells in the axial versus the appendicular skeleton may reflect either distinct roles for Foxc1 and Foxc2 in these two regions of the embryo; or a redundancy with other transcription factors that are restricted to the appendicular skeleton. Nevertheless, *Foxc* genes are important regulators of limb skeletogenesis, as *Prx1-cre;- Foxc1^Δ/Δ^;Foxc2^Δ/Δ^*embryos displayed smaller malformed fore- and hindlimbs. Limb defects appeared to be spatially patterned as formation of distal bone elements were more impacted compared to proximal ones. For instance, in the hindlimb, cartilage formation and bone mineralization was markedly reduced in the tarsals, metatarsals and phalanges of *Prx1-cre;Foxc1^Δ/Δ^;Foxc2^Δ/Δ^* embryos; while the tibia did form but was shorter and curved and the fibula was grossly truncated. In contrast, the femur did not appear noticeably different from control limbs. We propose that the proximal-distal patterning effects we observe are a result of a disruption of the temporal sequence of limb development rather than a spatial patterning effect. Limb structures develop in proximal to distal manner such that bones in the stylopod are specified and form before the zeugopod and autopod elements (Tickle and Towers, 2020). One hypothesis to account for the phenotypes we observe in *Prx1-cre;Foxc1^Δ/Δ^;Foxc2^Δ/Δ^* embryos is the possibility that fewer skeletogenic cells are available in such mutants, either through impaired differentiation of precursor cells into chondrocytes or reduced proliferation of the progenitor population; and thus this pool of cells is exhausted or diminished as distal structures form. Indeed, we have observed elevated expression of Foxc1 and Foxc2 mRNAs at the onset of condensation in the limb bud mesenchyme and also in immature chondrocytes ((Fig 2D-E; Almubarak et al., 2021) compared with a relatively reduced expression in more differentiated chondrocytes; thus suggesting that these transcription factors play a role in the early stages of chondrogenesis. Alternatively, the proximal-distal limb formation phenotypes we observe may reflect a temporal delay in Cre recombination activity in proximal tissues compared to distal ones. However, as proximal to distal differences in limb phenotype severity have also been observed in global *Foxc1* or *Foxc2* mutant embryos as well as in Col2-cre targeted mutants (Almubarak et al., 2021; Kume et al., 1998; Winnier et al., 1997), the proximal-distal differences in the limb skeleton following deletion of Foxc1 and Foxc2 may not be due to differential timing of Cre-mediated recombination. We also noted that *Prx1-cre;Foxc1^Δ/Δ^;Foxc2^Δ/Δ^* limbs exhibited reduced bone formation in the posterior zeugopod cartilage elements compared to the anterior zeugopod cartilage elements. In the forelimbs, the ulna was underdeveloped compared to the radius; and in the hindlimbs, the fibula was smaller than the tibia. Interestingly, fate-mapping of Msx1-expressing forelimb bud precursor cells has indicated that an influx of Msx1-expressing limb bud mesenchymal precursor cells into the posterior zeugopod cartilage element (i.e., the ulna) occurs prior to that of the anterior zeugopod cartilage element (i.e., the radius)(Markman et al., 2023). Perhaps this difference in the timing of the formation and differentiation of the anterior versus the posterior zeugopod cartilage elements renders the posterior zeugopod elements, with a more limited temporal window to support their formation, more sensitive to the absence of Foxc loss. We also observed that the growth of the autopod cartilage elements was severely decreased in *Prx1-cre;Foxc1^Δ/Δ^;Foxc2^Δ/Δ^*embryos, but that patterning of these elements was by and large normal. Recent single cell analysis of the developing mouse limb suggests that differentiation of skeletal elements in the limb occurs through a complex, three dimensional process and that different pools of progenitor cells give rise to specific elements (e.g. radius vs ulna; (Markman et al., 2023). Thus, the varying kinetics of precursor cell influx and cartilage differentiation of these differing cartilage elements may render these elements differentially affected by loss of Foxc1 and Foxc2.

The initial progression into chondrocyte hypertrophy was disrupted in *Prx1- cre;Foxc1^Δ/Δ^;Foxc2^Δ/Δ^* embryos. At E14.5, we observed a grossly smaller hypertrophic zone with lower levels of COLX production and reduced expression of *Ihh* in prehypertrophic chondrocytes. However, once chondrocytes become hypertrophic, growth plate defects in the *Prx1-cre;Foxc1^Δ/Δ^;Foxc2^Δ/Δ^*embryos were less evident. We propose that *Foxc1* and *Foxc2* function in the initial progression into chondrocyte hypertrophy, however once this differentiation state has occurred *Foxc* genes are not needed. It is well established that signaling molecules secreted by either prehypertrophic or hypertrophic chondrocytes regulate many aspects of chondrocyte differentiation and the endochondral ossification process. However, it’s important to consider that the first wave of chondrocyte differentiation occurs independently of hypertrophic chondrocytes, since these cells have yet to form. We propose that *Foxc1* and *Foxc2* function in the initial progression of chondrogenesis (hypertrophic-independent differentiation) and are dispensable for chondrocyte differentiation once hypertrophic chondrocytes are formed. This idea is supported by the reduction in *Foxc1* and *Foxc2* mRNA expression in more mature chondrocytes zones (Fig 2; (Almubarak et al., 2021). This notion of *Foxc1* and *Foxc2* functioning in the initial stages of chondrocyte generation from progenitor cells may also explain why loss of these factors blocks chondrocyte differentiation *in vitro* (Almubarak et al., 2021; Kume et al., 1998; Winnier et al., 1997; Yoshida et al., 2015).

The progression through or the exit from chondrocyte hypertrophy was delayed by loss of *Foxc1* and *Foxc2* function. Expanded hypertrophic chondrocytes zones and smaller POCs were observed from E16.5 onward in *Prx1-cre;Foxc1^Δ/Δ^;Foxc2^Δ/Δ^* embryos. COLX protein levels were expanded in the HC zone, yet *ColX* mRNA expression was not affected in this region in *Prx1-cre;Foxc1^Δ/Δ^;Foxc2^Δ/Δ^* embryos, suggesting that removal of COLX-positive cells was impaired. MMP13 localization and TRAP staining was also reduced in the POC of *Prx1- cre;Foxc1^Δ/Δ^;Foxc2^Δ/Δ^* mice, while no changes in TUNEL-signal were observed in this region, suggesting that the expansion of the hypertrophic chondrocyte zone was a result from impaired remodeling of the hypertrophic chondrocytes rather than changes in cell death.

Mineralization of the POC was affected in *Prx1-cre;Foxc1^Δ/Δ^;Foxc2^Δ/Δ^* embryos. Foxc1 and Foxc2 are required for osteoblast differentiation *in vitro* (Hopkins et al., 2016; Mirzayans et al., 2012; Park et al., 2011; Rice et al., 2003), however, impaired osteoblast differentiation alone likely does not account for the phenotypes we observe. In the perichondrium/periosteum surrounding the tibia in *Prx1-cre;Foxc1^Δ/Δ^;Foxc2^Δ/Δ^*, OSX-positive osteoblasts were present from E15.5 onward, although mineralization was delayed. In the presumptive POC of *Prx1- cre;Foxc1^Δ/Δ^;Foxc2^Δ/Δ^*embryos, OSX expressing osteoblasts formed but fewer were detected owing to the reduced size of the POC. Osteoblasts from the bone collar invade into the POC along with blood vessels. Vascularization of the POC did not seem to be affected as hypertrophic chondrocytes expressed *Vegfa* and blood vessels were present in the *Prx1-cre;Foxc1^Δ/Δ^;Foxc2^Δ/Δ^*POC (Gerber et al., 1999; Zelzer et al., 2004). However, as reduced OSX-labeled cells were detected in the POC of *Prx1-cre;Foxc1^Δ/Δ^;Foxc2^Δ/Δ^* embryos, it is possible that Foxc1-Foxc2 deficient osteoblasts are unable to associate with vascular endothelial cells when blood vessels populate the POC. We did observe persistent OPN localization and dramatically reduced expression of the *Phex* in the POC of *Prx1-cre;Foxc1^Δ/Δ^;Foxc2^Δ/Δ^* embryos. As PHEX acts to proteolytically process OPN for mineralization to proceed (Addison et al., 2010; Barros et al., 2013), loss of Phex expression in *Prx1-cre;Foxc1^Δ/Δ^;Foxc2^Δ/Δ^* leads to an accumulation OPN that disrupts mineralization in the POC. This reduced mineralization would lead to malleable bones and may cause the bent skeletal structures we observe in *Prx1-cre;Foxc1^Δ/Δ^;Foxc2^Δ/Δ^*mutant embryos. Whether FOXC1 and FOXC2 directly regulate *Phex* expression is a current research question that we are exploring.

In summary we report the overlapping roles for Foxc1 and Foxc2 in limb bud mesenchymal progenitors to regulate endochondral ossification in the appendicular skeleton (Fig 10). We propose that Foxc1 and Foxc2 function at two phases in endochondral ossification: during the initial differentiation of chondrocyte progenitors towards the formation of the hypertrophic zone, and later in the remodelling of hypertrophic chondrocytes to allow formation of the POC and marrow space.

**Fig. 10.**
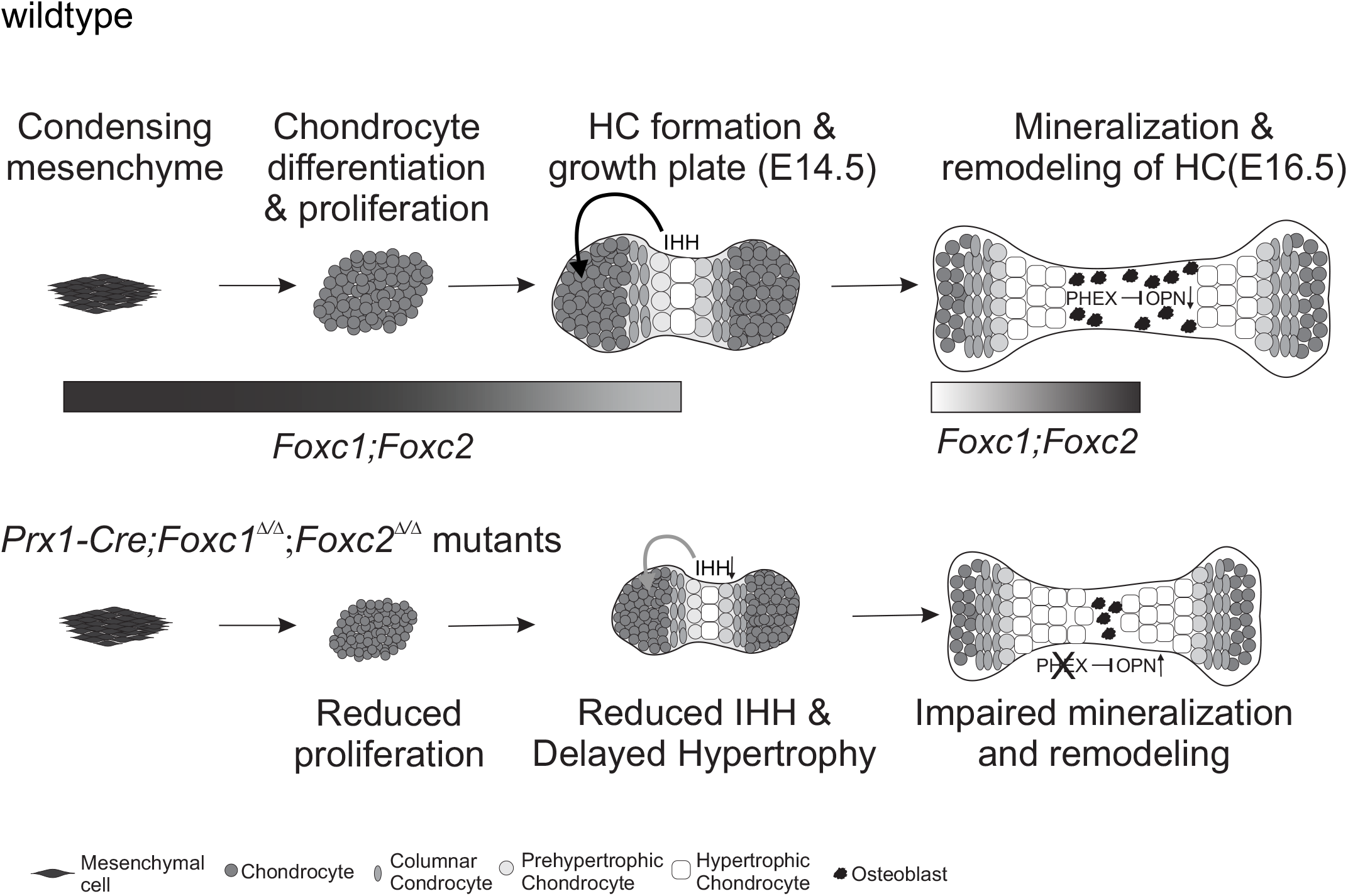
Foxc1 and Foxc2 function at two separate phases in endochondral ossification of the limb. We propose that *Foxc1* and *Foxc2* act at different stages of endochondral ossification of the limb skeleton (depicted by shaded bar). First, these factors function in the differentiation of limb bud mesenchyme towards the chondrocyte lineage. Here, *Foxc1* and *Foxc2* regulate chondrocyte proliferation and the initial formation of hypertrophic chondrocytes. In *Prx1- cre;Foxc1^Δ/Δ^;Foxc2^Δ/Δ^*mutants we observe a delayed entry into hypertrophy with fewer *Ihh*- expressing pre-hypertrophic chondrocytes and hypertrophic chondrocytes present. Second, once hypertrophic chondrocytes have formed and the growth plate organized (after E14.5), *Foxc1* and *Foxc2* function in growth plate chondrocytes is not as crucial. *Foxc1* and *Foxc2* function is required to remodel and remover the hypertrophic chondrocytes to create the primary ossification centre. At this stage Foxc1 and Foxc2 are required for *Phex* expression which functions process OPN and allow osteoblast mineralization to occur.

## Materials and Methods

### Mouse models

Experiments using mouse models were either approved by the University of Alberta Animal Care and Use Committee (AUP804) or approved by the Harvard Medical School Institutional Animal Care and Use Committee (IACUC). To explore whether Foxc1 and Foxc2 share overlapping roles in Sox9 expressing cells, *Foxc1^fl/fl^ ;Foxc2 ^fl/fl^* (Sasman et al., 2012) mice were with mated mice containing *Sox9^ires-Cre/+^*(Akiyama et al., 2005) to generate E12.5 embryos which either deleted neither, one or both of these Foxc family members.

*Prx1-cre;Foxc1^Δ/Δ^;Foxc2^Δ/Δ^*mince were generated through crossing *Foxc1^fl/fl^ ;Foxc2^fl/f^*^l^ (Sasman et al., 2012) with Prx1-cre ^+/-^ mice (Logan et al., 2002). Timed pregnancies were performed by crossing male *Prx1-cre ^+/-^ ;Foxc1 ^+/fl^ ;Foxc2 ^+/fl^* mice to female *Foxc1 ^fl/fl^ ;Foxc2 ^fl/fl^* mice. The day of the detection of a vaginal plug was denoted as E0.5. We genotyped weaned mice using ear notch biopsies and embryos using skin. The genotyping process was conducted using the KAPA mouse genotyping kit (Millipore Sigma) using the following primer pairs: *Foxc1* (forward 5’- ATTTTTTTTCCCCCTACAGCG-3’; reverse 5’-ATCTGTTA GTATCTCCGGGTA-3’), *Foxc2* (forward 5’ CTCCTTTGCGTTTCCAGTGA -3’; reverse 5’- ATTGGTCCTTCGTCT TCGCT - 3’) and *Prx1-cre* (forward 50 -GCCTGCATTACCGGTCGATGCAACGA-30; reverse 50 - GTGGCAGATGGCGCGGCAACACCATT-30). All experimental comparisons were made between littermates, with a minimum of three litters used per experiment, unless otherwise noted.

### Whole Skeleton Staining

Embryos were collected at E12.5, E15.5 and E18.5 and processed for whole skeleton Alcian blue and alizarin red staining as described in Rigueur and Lyons (2014).

### Tissue preparation

Tissues were dissected at specific stages and fixed in 4% paraformaldehyde at 4°C overnight before embedding in paraffin. Sections were cut at 5 μm thickness and collected on Superfrost- plus slides (Fisher Scientific). E17.5 limbs were decalcified with EDTA at 4°C overnight before paraffin embedding.

### Histology

All sections were first dewaxed with xylene and rehydrated with graded ethanol and water. For Safranin O staining sections were stained with hematoxylin for 8 minutes and rinsed with running tap water for 10 minutes. Next, sections were stained with 0.001% Fast green for five minutes, followed with 1% Acetic acid wash for 10-15 seconds to stabilize the staining. Slides were then stained with 0.1% Safranin O for five minutes, rehydrated with 100% ethanol and xylene, and mounted with coverslips. For Alcian blue Von-kossa staining, sections were incubated with 1% silver nitrate solution under ultraviolet (UV) light for 20 minutes. Slides were then rinsed with two water changes followed by five minutes incubation with 5% sodium thiosulfate to remove the unreacted silver. Sections were then stained with Alcian blue for 30 minutes and counterstained with nuclear fast red for five minutes. For TRAP staining, sections were then incubated in a pre-warmed TRAP Staining solution Mix at 37°C for 30 minutes. Next, slides were rinsed with water and counterstained with 0.02% Fast green.

### *In situ* hybridization

Fluorescent ISH for multiplex was performed using RNA scope Multiplex Fluorescent kit following the manufacturer’s protocol for paraffin embedded sections (Advanced Cell Diagnostics). Sections were boiled in antigen retrieval solutions for 15 minutes. The following probes were used. Negative control (REF: 310043); *Foxc1* (REF: 412851); *Foxc2* (REF: 406011); *Fgfr3* (REF: 440771); *Fgfr1*(REF: 454941); *Ihh* (REF: 413091); *Gli1*(REF:311001) ; *Gli2* (FEF: 405771-C2); *Gli3* (REF:445511); *Pthlh* (REF: 456521); *Pth1r* (REF: 426191) ; *Ptch1* (REF: 402811); *Ptch2* (REF: 435131); *Colx* (REF:433491); *Tnfsf11*(REF: 410921); *Vegf* (REF: 436961); and *Phex* (REF: 426201).

### Immunofluorescence

Paraffin sections were collected and slides prepared as described above. For SOX9, OSX, OPN, KI67, and RUNX2 antibodies, antigen retrieval was performed through boiling the slides in citrate buffer (10 mM trisodium citrate, pH 6.0; 0.05% Tween20) for 20 minutes. For COL1, COL2a, COLX, MMP13, VEGFA antibodies, samples were incubated in hyaluronidase for 30 minutes at 37°C. Next, slides were blocked in 5% donkey serum in PBS with 0.05% Triton X- 100 (PBSX) for one hour. Slides were then incubated with the primary antibody overnight at 4°C. The following antibodies were used for immunofluorescence microscopy: COL IIa (Abcam,ab185430, 1:100); COLX (Abcam, ab5832, 1:50); COL I (Abcam,ab88147; GR3225500-1, 1:100); MMP13 (Abcam, ab39012, 1:100); RUNX2 (Abcam,ab76956, 1:200), IB4 (Thermo Fisher Scientific cat: VECTB1205, 1:500), SOX9 (MilliporeSigma, AB55535, 1:200); OPN (SCBT, sc22536-R, 1:100); OSX (SCBT, sc21742, 1:100); KI67 (Bethyl, IHC00075, 1:100).

### TUNEL (terminal deoxynucleotidyl transferase dUTP nick end labeling) assay

Apoptosis was detected using the *In situ* Cell Death Detection Kit, TMR red (Roche). Tibia sections were obtained and processed as described above. Slides were permeabilized with proteinase K working solution (10μg/ml in 10mM Tris/HCL, PH 7.4-8) for 30 minutes at 37°C and washed twice with 1xPBS. Then, sections were treated with the TUNEL reaction mixture for 60 minutes in a humidified atmosphere at 37°C. Slides were then washed three times with 1xPBS and stained with DAPI for 5 minutes and mounted with Prolong Gold anti-fade reagent (Invitrogen).

## Acknowledgements

We would like to thank Dr Karen Lyons for helpful comments on the manuscript.

## Data Availability

All data generated in this study are provided in this manuscript.

## Supplementary Figures

**Fig. S1.**
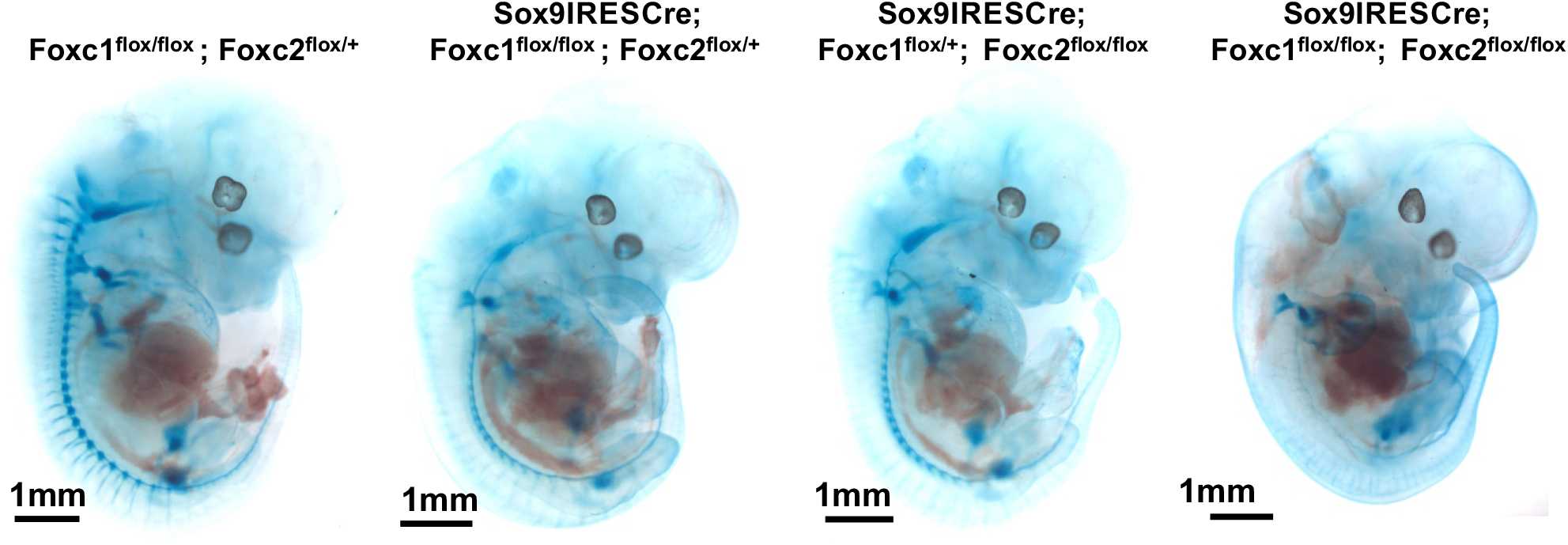
*Foxc1* and *Foxc2* share redundant roles to promote axial chondrogenesis. Whole embryos littermates embryos at E12.5 were stained with Alcianblue to detect chondrogenic differentiation. Similar results have been obtained with embryos from 2 litters, containing 3 embryos of the least frequently occurring genotype (i.e., *Sox9^ires-Cre/+^;Foxc1^flox/flox^;Foxc2^flox/flox^*).

**Fig. S2.**
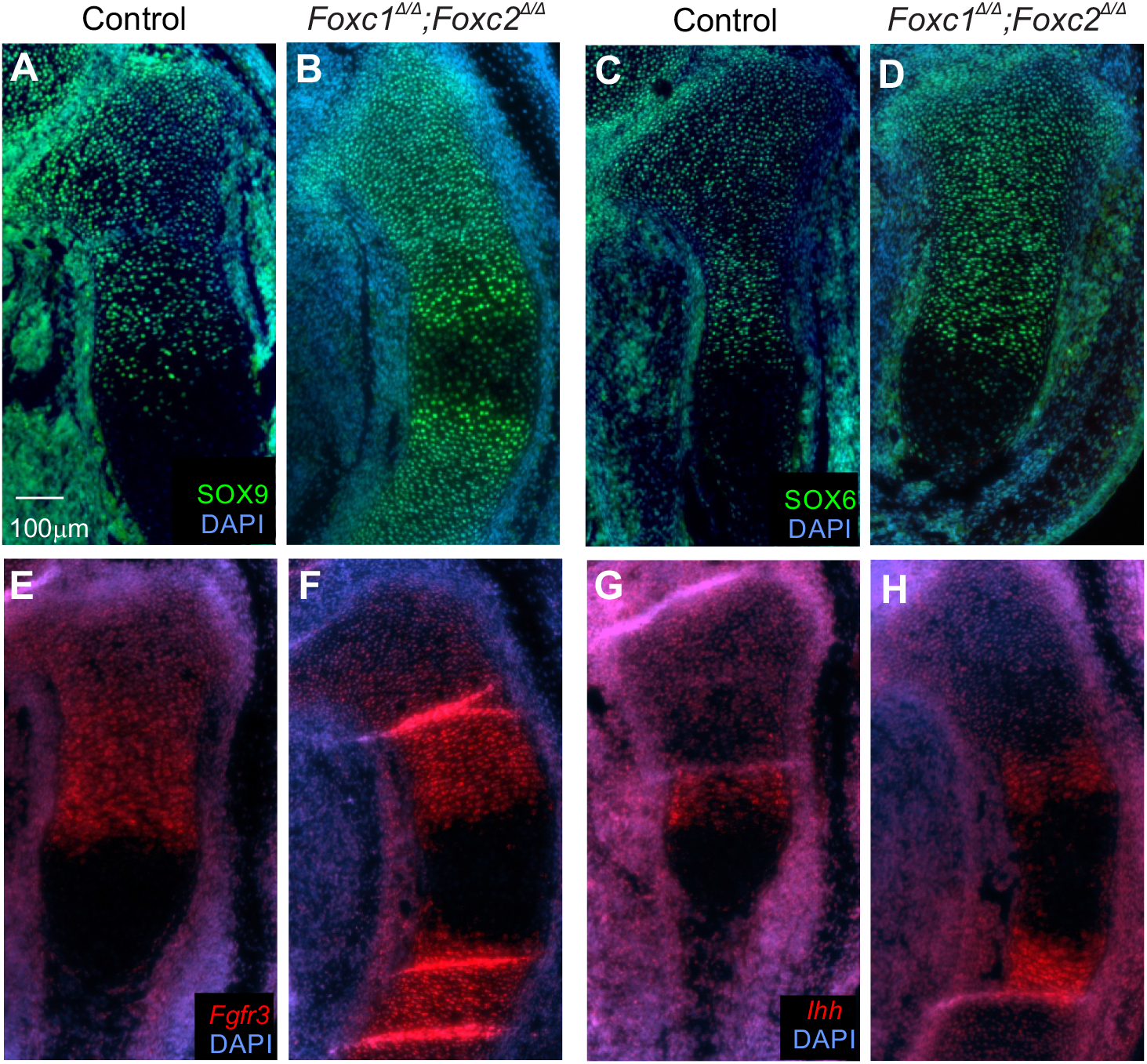
Growth plate formation is not affected by loss of Foxc1 and Foxc2. Chondrocyte differentiation and growth plate formation in the proximal tibia at E15.5 was examined. Immunofluorescence analysis of SOX9 (A, B), SOX6 (C, D) in the the resting zone and columnar chondrocytes in control and *Prx1-cre;Foxc1^Δ/Δ^;Foxc2^Δ/Δ^*embryos. *Fgfr3* (E, F) and *Ihh* (G, H) mRNA expression was visualized by in situ hybridization to demarcate the proliferating chondrocytes and the pre-hypertrophic chondrocytes, respectively. Scale bar 100 μm.

**Fig. S3.**
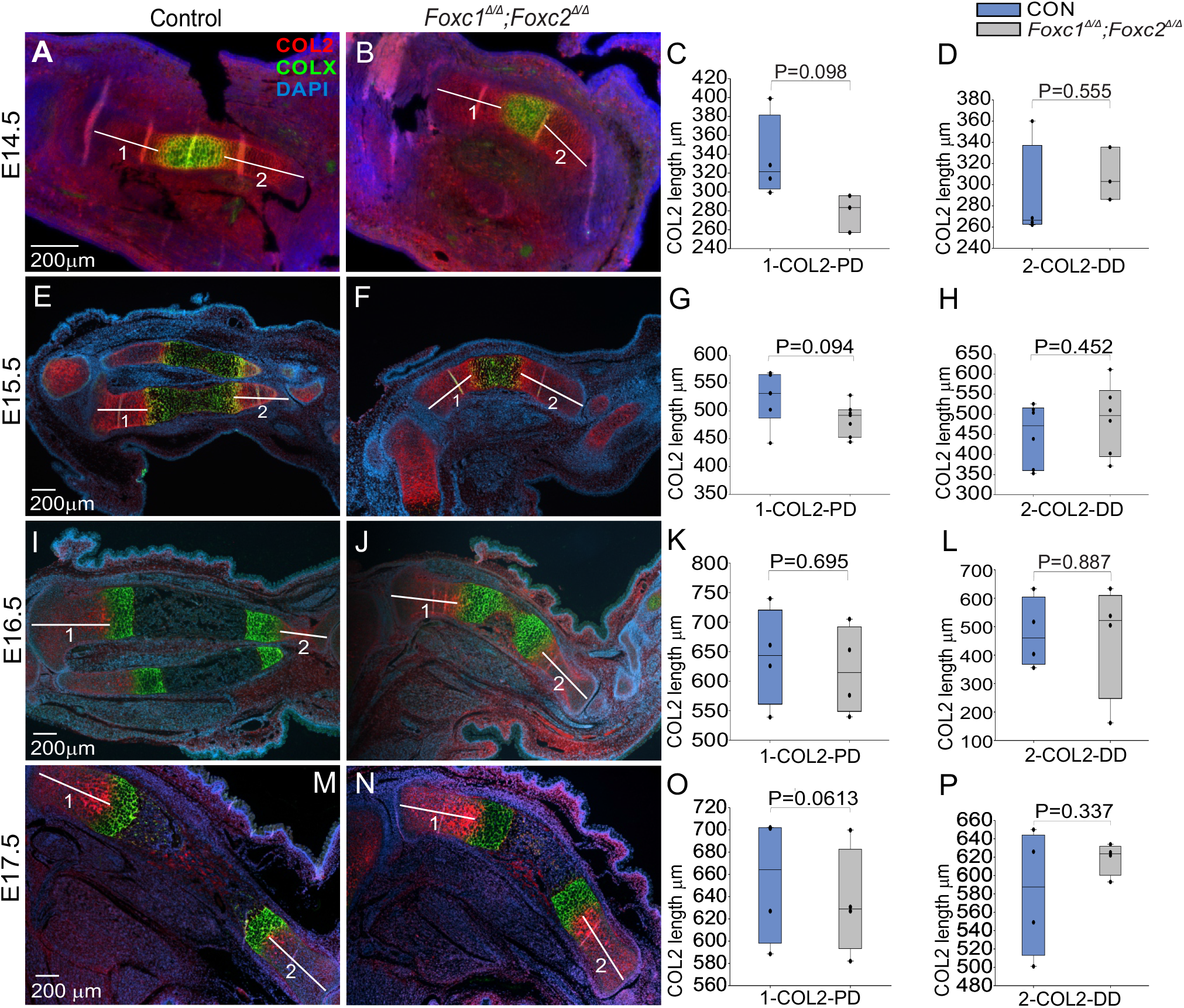
Length of COL2-expressing chondrocyte zone is unaffected by deletion of *Foxc1* and *Foxc2*. Comparison of the COL2 (red) and COLX (green) chondrocytes in control and *Prx1- cre;Foxc1^Δ/Δ^;Foxc2^Δ/Δ^* tibia. The length of the COL2 (red) chondrocytes was measured from the distal and proximal tibia at E14.5 (A-D), E15.5 (E-H), E16.5 (I-L) and E17.5 (M-P). The length of COL2 positive chondrocyte zone did not significantly change. Scale bar 200 μm. Statistical analysis was performed using Student t-test. (n=4).

**Fig. S4.**
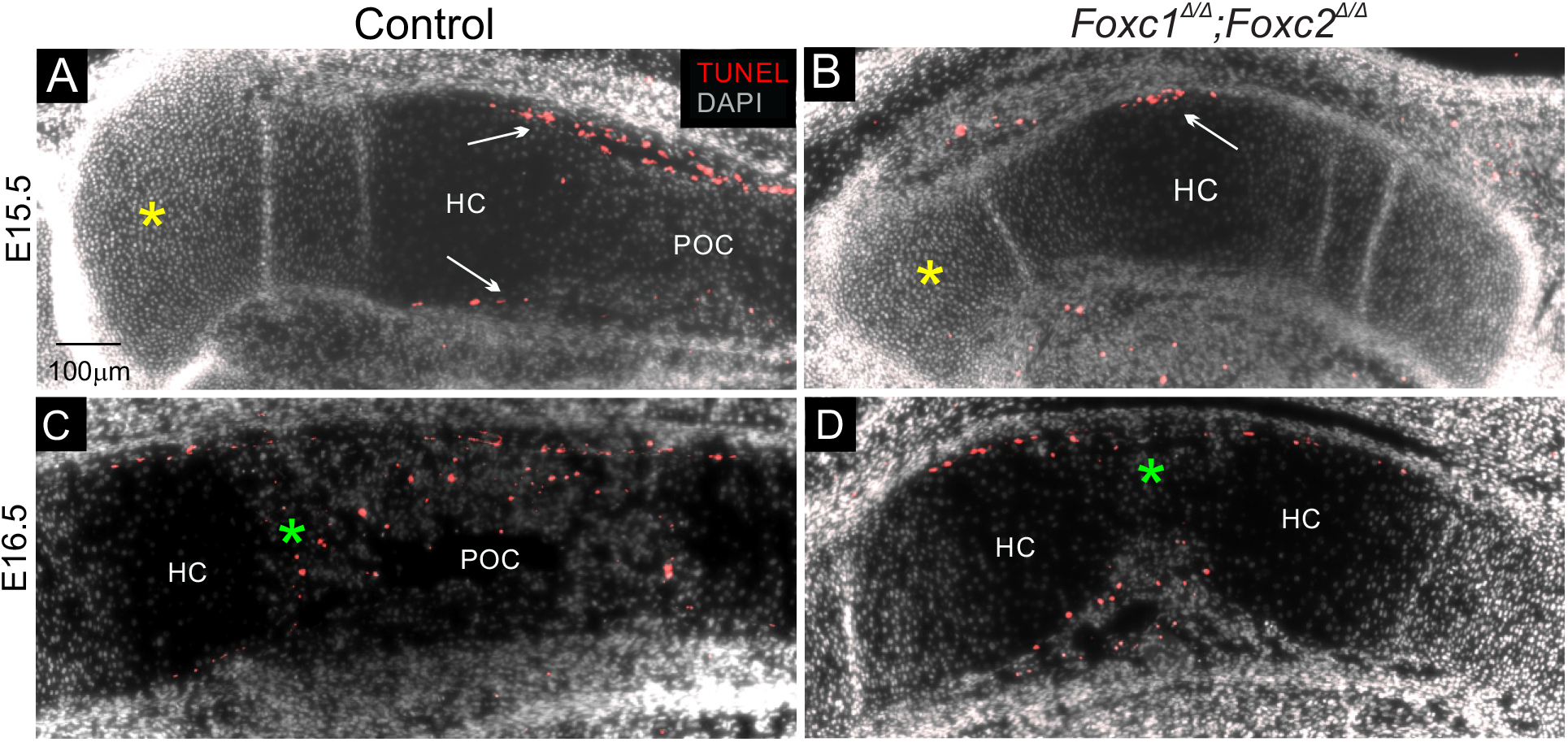
Expanded hypertrophic chondrocyte zone is not a result of decreased cell death in the growth plate of *Prx1-cre;Foxc1^Δ/Δ^;Foxc2^Δ/Δ^* limbs. Cell death was monitored by TUNEL assays. was performed in E15.5 and E16.5 control and *Prx1-cre;Foxc1^Δ/Δ^;Foxc2^Δ/Δ^* tibia. Distinctive apoptosis activity in a cell population adjacent to the perichondrium and the periosteum (white arrows) was observed at E15.5. However, no cell death was detected in the growth plate chondrocytes in both control and *Prx1-cre* mutant limbs (yellow asterisk) (A, B). At E16.5 control limbs displayed some TUNEL signal in the POC (green asterisk; C) . Reduced TUNEL signal was detected within the smaller POC in the *Prx1- cre;Foxc1^Δ/Δ^;Foxc2^Δ/Δ^* embryos (green asterisk; D). (n=3)

